# High-yield Production of Recombinant Platelet Factor 4 Protein by Harnessing and Honing the Gram-negative Bacterial Secretory Apparatus

**DOI:** 10.1101/830851

**Authors:** Saeed Ataei, Mohammad Naser Taheri, Fatemeh Taheri, Farahnaz Zare, Niloofar Amirian, Abbas Behzad-Behbahani, Amir Rahimi, Gholamhossein Tamaddon

## Abstract

**Background:** Platelet factor 4 is a cytokine released into the bloodstream by activated platelets and plays a pivotal role in heparin-induced thrombocytopenia etiology and diagnosis. Therefore, a sustainable source of recombinant PF4 with structural and functional similarity to its native form is urgently needed to be used in diagnostic procedures.

To this end, a three-in-one primary construct was designed and custom synthesized based on the pET26b backbone from which three secondary constructs could be derived each capable of employing either type I, type II secretory or cytoplasmic pathways. Protein expression and secretion were performed in *Escherichia coli* BL-21 (DE3) and were confirmed by SDS-PAGE and Western blotting. To further enhance protein secretion, the effect of several controllable factors including IPTG, Triton X-100, Sucrose, and Glycine were individually investigated at first. In the next step, according to fractional factorial approach, the synergistic effect of IPTG, Triton X-100, and Glycine on secretion was further investigated. To ascertain the structure and function of the secreted recombinant proteins, Dynamic light scattering was utilized and confirmed rPF4 tetramerization and heparin-mediated ultra-large complex formation. Moreover, Raman spectroscopy was exploited to determine the rPF4 secondary structure.

**Results:** Type II secretory pathway was proven to be superior over type I in case of rPF4 secretion into the extracellular milieu. Protein secretion mediated by Type II was enhanced to approximately more than 700 μg/ml. Large quantities of native rPF4 up to 20 mg was purified upon a minor scale up to 40 ml of culture medium. Dynamic light scattering unveiled native rPF4 quaternary structure revealing the formation of tetramers having an average size of 10 nm and formation of larger complexes of approximately 100-1200 nm in size following heparin supplementation, implying proper protein folding, tetramerization, and antigenicity. Analysis of the Zeta potential on approximately 600 μg/ml of rPF4 revealed a 98 mV positive charge which further confirms protein folding. Moreover, rPF4 secondary structure was determined to be 43.5% Random coil, 32.5% β-sheet, 18.6 % α-helix and 4.9 % Turn, which is in perfect agreement with the native structure.

**Conclusion:** our results indicate that the gram-negative type II bacterial secretory system holds a great promise to be employed as a reliable protein production strategy with favorable industrial application. However, further efforts are required to realize the full potential of secretory pathways regarding their application to proteins with distinct characteristics.

Graphical Abstract.
rPF4 secretion mediated by type 2 secretory system. The pelB signal sequence directs protein export into the extracellular milieu through the SecYEG translocon complex in a process assisted by SecB chaperone. A) Indicates protein secretion before supplementation with additives and B) indicates secretion in the presence of additives.

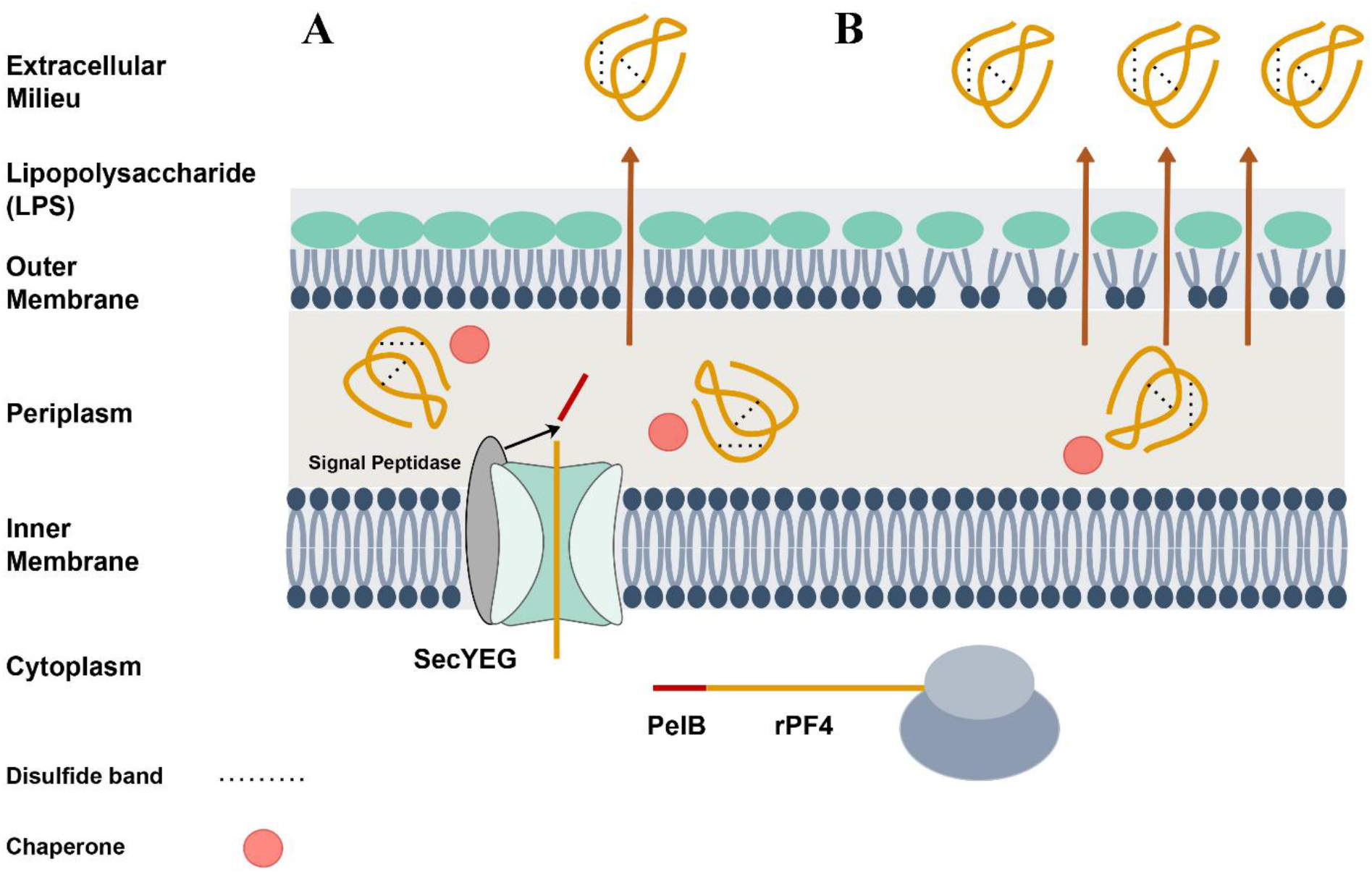

## 1. Background

Heparin-induced thrombocytopenia (HIT) is a deleterious drug reaction caused by heparin administration in which platelet factor 4 (PF4) as a positively charged protein binds heparin, and the resulting PF4-Heparin complex adversely stimulates an immune response. Following engagement with the FcγIIa receptors, the PF4-Heparin-IgG complex activates platelets and therefore gives rise to thrombosis. The disease etiology also includes antibody-mediated endothelial trauma or excessive tissue factor production in case the antigen-antibody complexes interact with monocytes (1–5). Currently, there are several approaches to diagnosing patients with HIT, with some of them requiring exogenous PF4(6, 7). Therefore, a cost-effective, sustainable, and scalable source of recombinant PF4 with antigenic and biological characteristics similar to the native platelet-derived PF4 is imperative.

PF4 is a 7.8 kDa protein, consisting of 70 amino acids in its mature form and possesses two disulfide bonds. (8) PF4 is released during platelet activation and plays a plethora of biological processes including regulation of angiogenesis, megakaryopoiesis, and activation or proliferation of leukocytes (9–12).

Gram-negative bacteria are known to have at least six distinct protein secretory pathways. Type I and II have been mainly used for recombinant protein production. (13)

Type I secretion system (T1SS) protein substrates are characterized by a non-cleavable c-terminal signal sequence recognized by the components of the T1SS machinery and are directly secreted to the extracellular milieu. T1SS protein machinery is composed of HlyB, an ATP-binding cassette (ABC) transporter, and HlyD, a protein that connects the inner membrane to the outer membrane. The T1SS system is epitomized by the HylA protein of pathogenic *E. coli* (14–16).

Type II secretion system (T2SS) has been found to be a well-suited pathway for the production of some recombinant proteins. In this system, substrates are characterized by an N-terminal and cleavable signal sequence. Protein folding takes place following signal peptide cleavage and protein translocation into the periplasmic compartment(13, 16). Numerous recombinant proteins have been successfully produced by utilizing the aforementioned system including Thermobifida fusca cutinase(17, 18), Phospholipase D(19), pullulanase(20) and archeal lipase(21). Many factors such as protein stability, function, immunogenicity and proper folding, etc. need to be considered when it comes to producing a recombinant protein. Unlike the cytoplasmic protein expression, the extracellular protein secretion meets many of the above-mentioned requirements for the production of proteins. Protein secretion using type I and type II systems seems to be amenable to efficient and functional production of rPF4, being small in size, bearing two disulfide bonds, and urgently needed to be produced in large scales. Therefore, a three-in-one system was designed enabling cytoplasmic protein expression or secretion from type I or type II systems through derivative constructs, allowing us to evaluate the potentiality of each system for large scale and functional production of rPF4 protein.

## 2. Results

### 2.1 Construction of rPF4, pelB-rPF4 and rPF4-HLAs

The recombinant pelB-rPF4, rPF4-HLAs (312 bp and 471 bp respectively) (Fig. 2 and 3) and the rPF4 (264 bp) (Fig. 4) constructs, were derived from the three-in-one construct, ensuing digestions by NdeI, SalI and SlaI as described in section 5.2. The correct vector constructions were ultimately confirmed by colony PCR and sequencing

**Fig 1.**
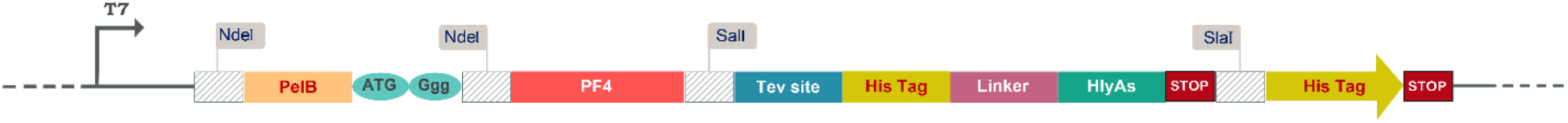
Schematic representation of the three-in-one construct design. pelB and HLAs signal sequences were placed upstream and downstream of the PF4 coding sequence respectively. The PF4 gene was fused to HlyAs through a linker sequence bearing a Tev recognition site.

**Fig 2.**
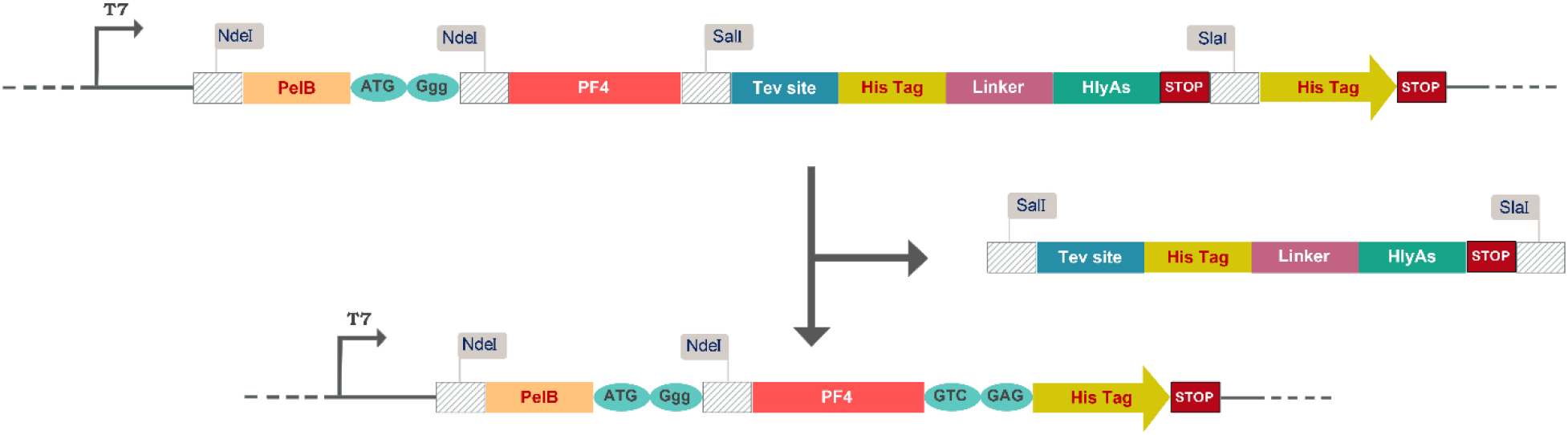
Schematic representation of the T2SS construct. By sequential digestions using SalI and SlaI restriction enzymes the fragment harboring HLAs and a His-tag is removed, as the part of the vector, and the His-tag along with the adjacent stop codon is placed in frame with PF4 coding sequence, converting the primary three-in-one system in to T2SS construct.

**Fig 3.**
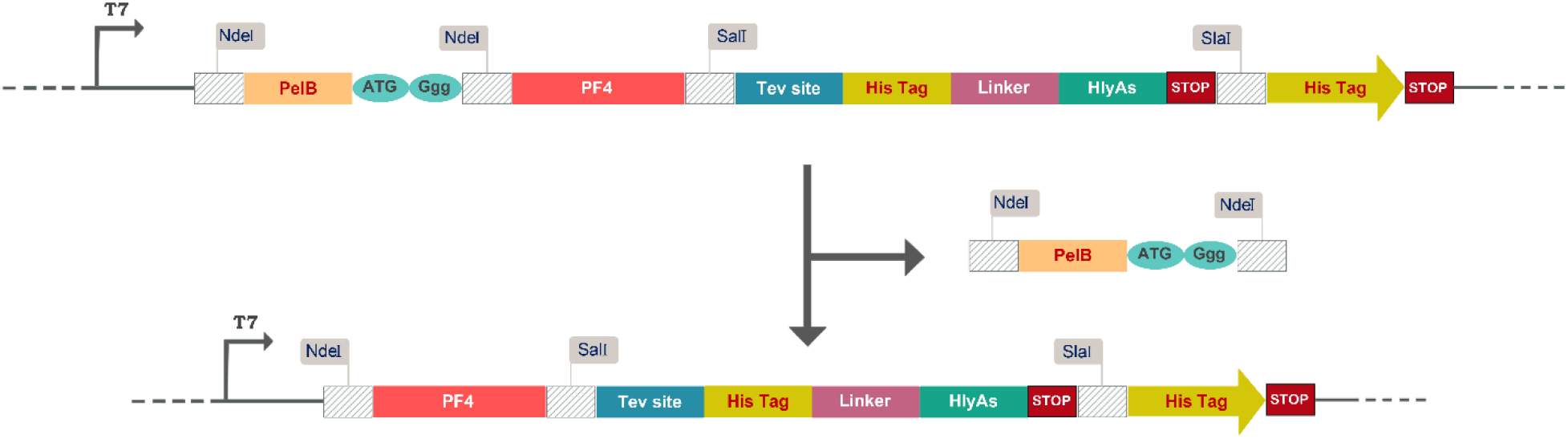
Schematic representation of the T1SS construct. As the pET26b vector harbors a NdeI restriction site upstream of the pelB leader peptide, another NdeI site was placed at the 5’ end of the PF4 coding sequence; as a result, through a single digestion using NdeI, the fragment bearing the pelB sequence is excised out and the T2SS construct is generated.

**Fig 4.**
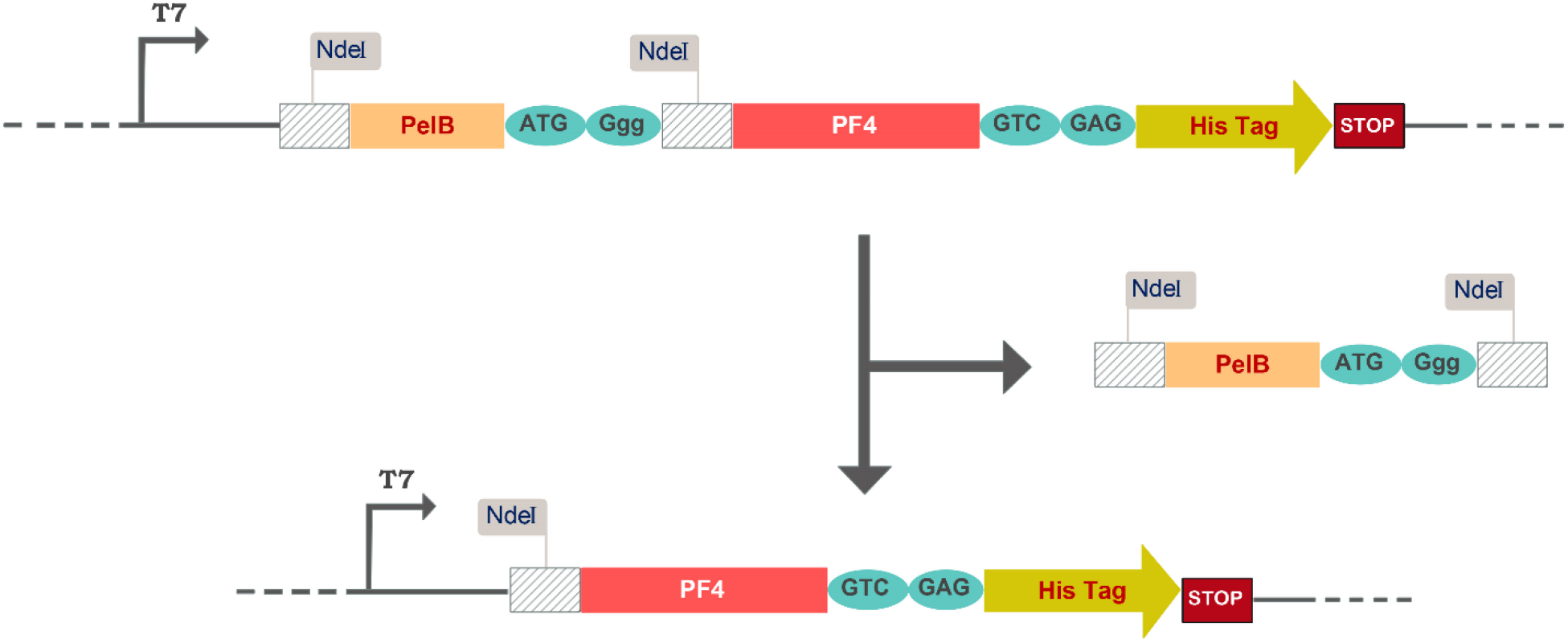
Schematic representation of the construct a enabling the cytoplasmic protein expression. The T2SS construct is subjected to NdeI digestion resulting a construct devoid of pelB and HLAs sequences.

### 2.2 Recombinant rPF4, pelB-rPF4 and rPF4-HLAs proteins expression and verification by SDS-PAGE

Protein expression was triggered by the addition of IPTG to the final concentration of 2 mM as described in section 5.3 and later the bacterial cleared lysate was subjected to SDS-PAGE analysis. Protein expression is triggered at a higher concentration of IPTG, results in protein aggregation and inclusion body formation which is not amenable to proteins SDS-PAGE analysis. This conundrum was solved by taking advantage of 8 M urea to disentangle the inclusion body formations. Following sonication, the protein bands of interest corresponding to the rPF4, pelB-rPF4, and rPF4-HLAs with respective molecular weights of 8.8, 11.4, and 16.2 kDa were detected in the soluble phase of cleared lysate (Fig. 5)

**Fig 5.**
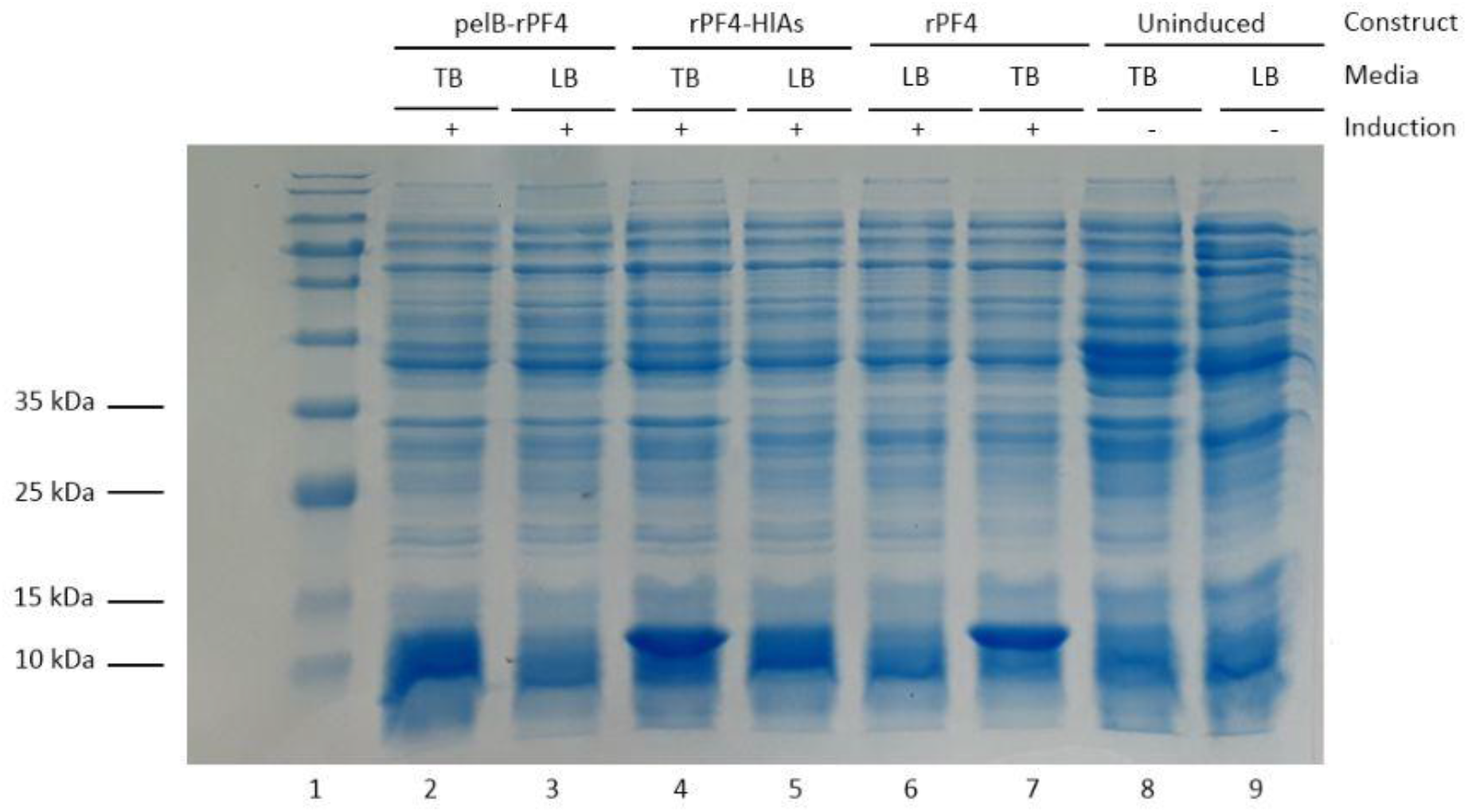
Analyzing cytoplasmic expression of the T1SS, T2SS and the cytoplasmic rPF4 constructs. Expression was induced by 2 mM IPTG for 2 hours in LB or TB media. Bacterial cleared lysates were subjected to analysis using SDS-PAGE. All the constructs indicated an efficient cytoplasmic protein expression, however, the TB media as opposed to LB, significantly improved the protein expression. As an inherent feature of the Lac operon, a minor leaky expression of rPF4 was observed in the non-induced groups.

### 2.3 Recombinant rPF4 protein purification

To confirm the cytoplasmic expression of rPF4, by use of the Ni-NTA matrix, the denaturing column protein purification method was performed. SDS-PAGE analysis on a 13.5% gel revealed that the first elution harbors the bulk of the desired proteins corresponding to approximately 7.8 kDa of weight as outlined in Fig. 6

**Fig 6.**
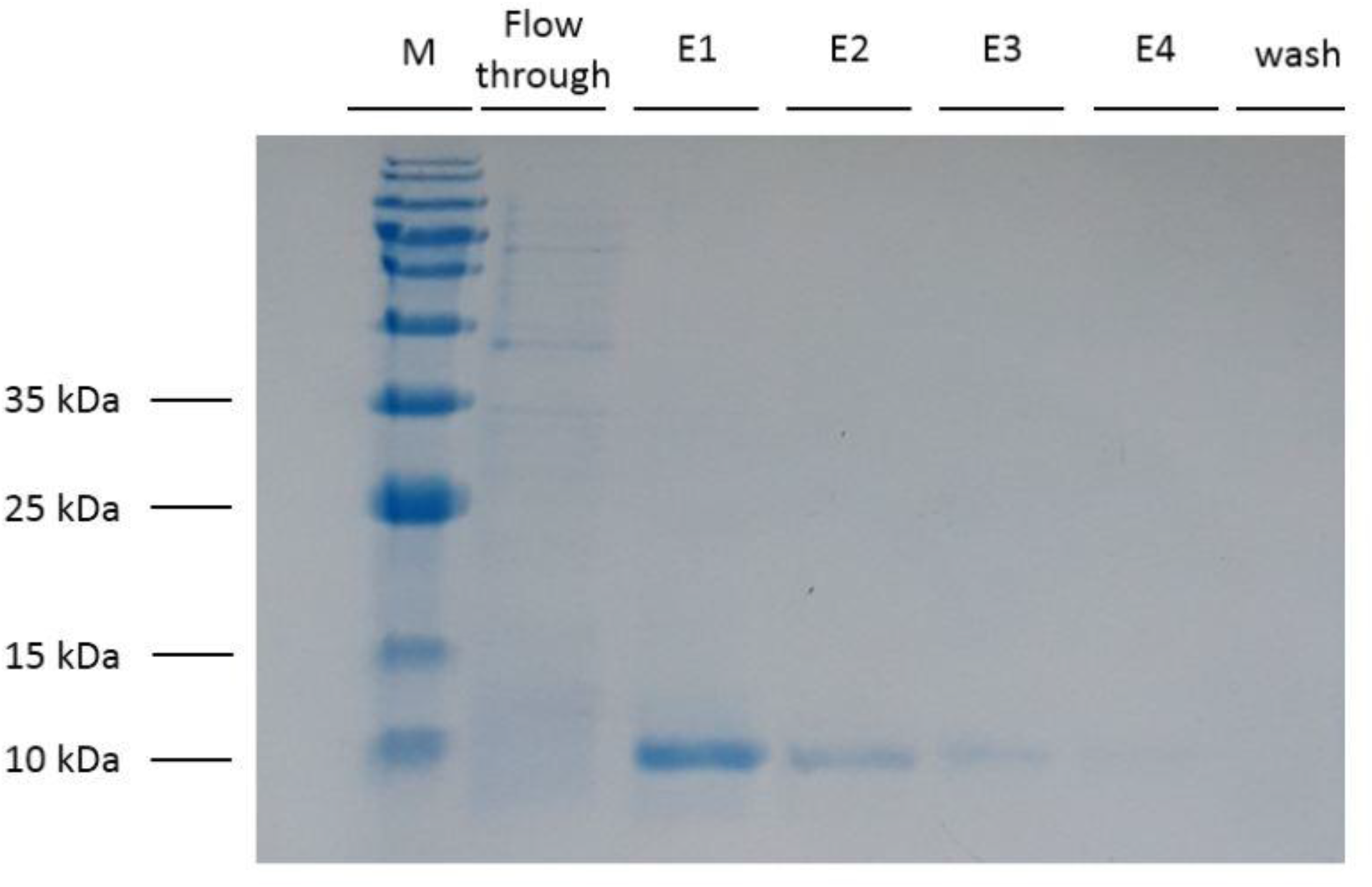
Purification of the recombinant rPF4 protein produced by the cytoplasmic expression system. Denaturing batch protein purification was performed. E1 to E4 correspond to 0.5-mL elution of the captured proteins. To alleviate contamination with unwanted proteins, the matrix was washed with the washing solution containing 10 mM of imidazole. M denotes the molecular weight marker

### 2.4 Recombinant rPF4, pelB-rPF4 and rPF4-HLAs proteins secretion and verification by SDS-PAGE and Western blotting

Mild IPTG induction and lower growth temperature helped with the protein folding process and further avoided inclusion body formation which is an important point for protein secretion. The bacterial cells were pelleted and the aqueous media from the cells was separated. Subsequently, a protein purification on TB and LB aqueous media was performed and the protein secretion into the extracellular milieu was analyzed by SDS-PAGE (Fig 7).The hosts bearing the pET26b-rPF4 construct were not able to secrete any protein into the extracellular environment. As depicted in figure 7, T2SS exerted a higher efficiency than T1SS in secreting rPF4 into the extracellular milieu, as a result, the T2SS was elected as the main strategy to carry on for the future endeavors. Furthermore, as evidenced by SDS-PAGE, protein expression and secretion were greatly enhanced in TB compared to LB media. Ensuing SDS-PAGE analysis, the proteins were blotted onto the PVDF membrane and the secreted rPF4 was confirmed by Western blotting (Fig. 8)

**Fig 7.**
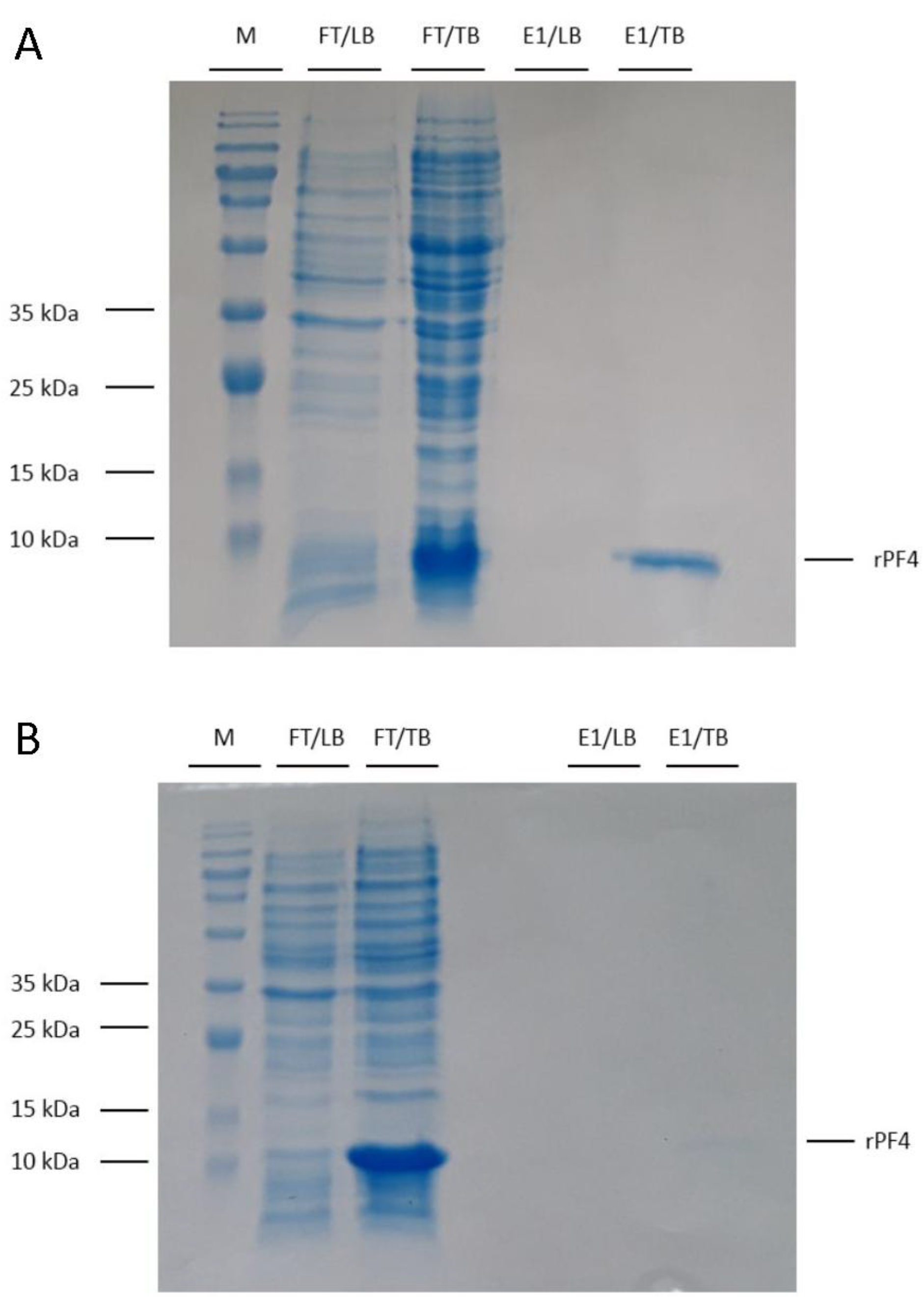
Analyzing the T1SS and the T2SS secretion systems. Protein expression and secretion were carried out in 7 mL of LB or TB media for 18 hours. Induction was triggered with 0.5 mM IPTG. FT refers to the flow through collected after the media passed through the column. E correspond to 0.5 mL fractions collected following elution of the proteins captured on the column. (A) Represents the T2SS mediated rPF4 protein secretion into the extracellular milieu using pelB leader peptide. (B) Represents the T1SS mediated rPF4 protein secretion into the extracellular milieu using HLA signal peptide. As the figures clearly depict, the T2SS is more efficient for secreting rPF4 than T1SS. Furthermore, as confirmed by the SDS-PAGE analysis growth in TB medium explicitly enhances the protein secretion process and is quite superior over LB medium.

**Fig 8.**
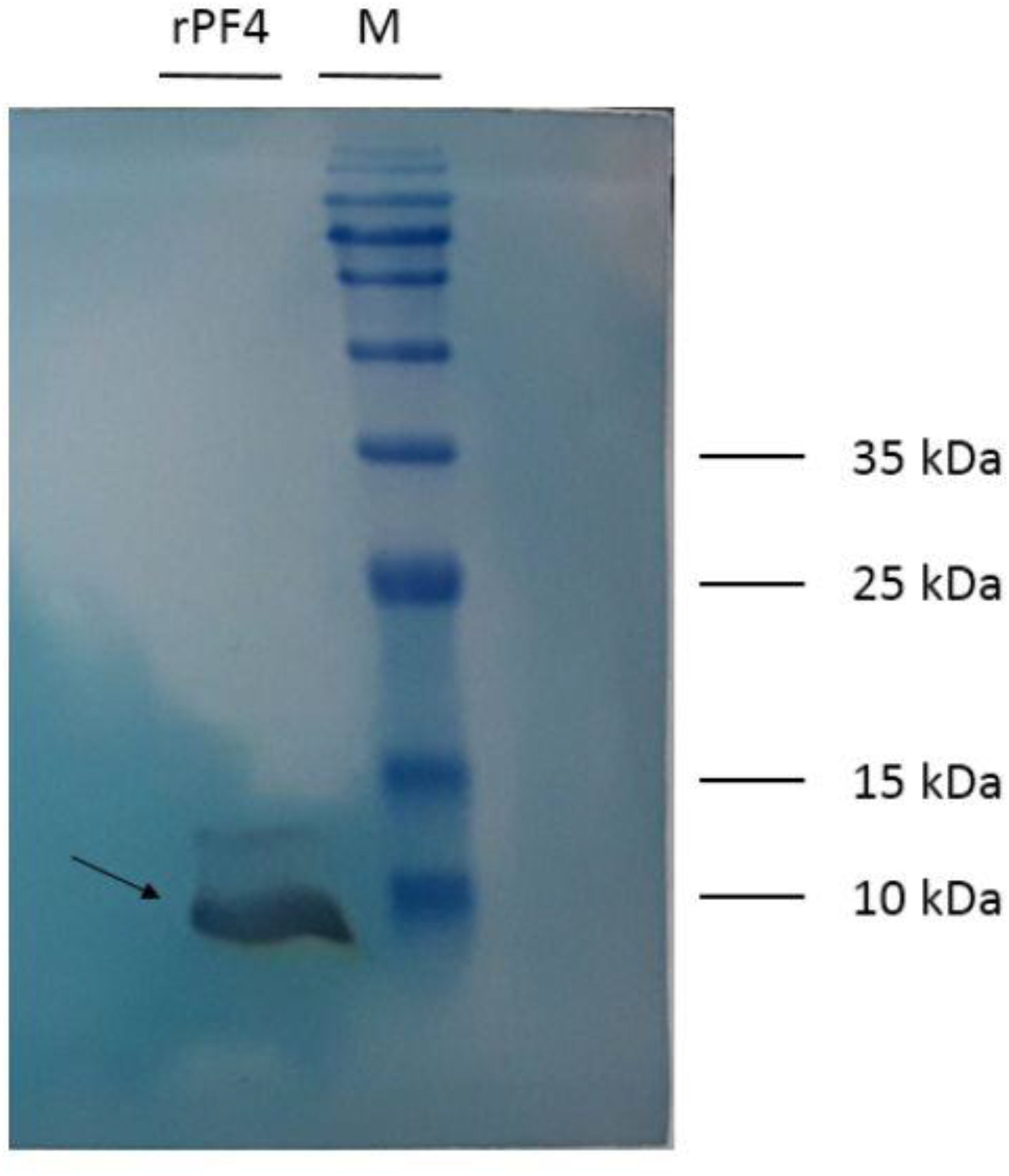
Western blotting confirmation of the rPF4 secretion into the outside medium. The T2SSmediated secreted rPF4 was collected from the extracellular milieu and was transferred from the gel onto the PVDF membrane. Following the treatment with a specific antibody the PF4 protein band was visualized. The arrow indicates the PF4 protein band. M denotes the molecular marker.

### 2.5 Individual effects of IPTG, Triton X-100, glycine, and sucrose on protein secretion rate

Initially the effect of various amounts of glycine (0%, 0.5%, 1%, 1.5%, 2%, 3%, 4%, 5% w/v), was analyzed. The level of protein secretions was augmented with the addition of 0.5%, glycine to the culture medium (Fig.9a) Further supplementation of 3%, 4%, and 5% glycine led to cellular demise as no cultural growth of bacteria was observed. Conversely, analyzing 5% and 10% sucrose supplementation subjected a negative effect on rPF4 release into extracellular milieu (Fig.9b). Moreover, the commencement of Triton X-100 was investigated, nevertheless, Bacterial cells succumbed to death by supplementation of 1%, 2% and 3% Triton X-100 as no cultural growth in the media was noted. Furthermore, the effect of rising concentrations of 0.01, 0.05, 0.1, 0.2, 0.3, 0.4, 0.5 mM IPTG on protein secretion was explored. Protein secretion was enhanced via 0.01, 0.05, 0.1 and 0.2 mM commencement of IPTG, however, higher concentrations of IPTG negatively affected protein secretion rate (Fig.9c).

**Fig 9.**
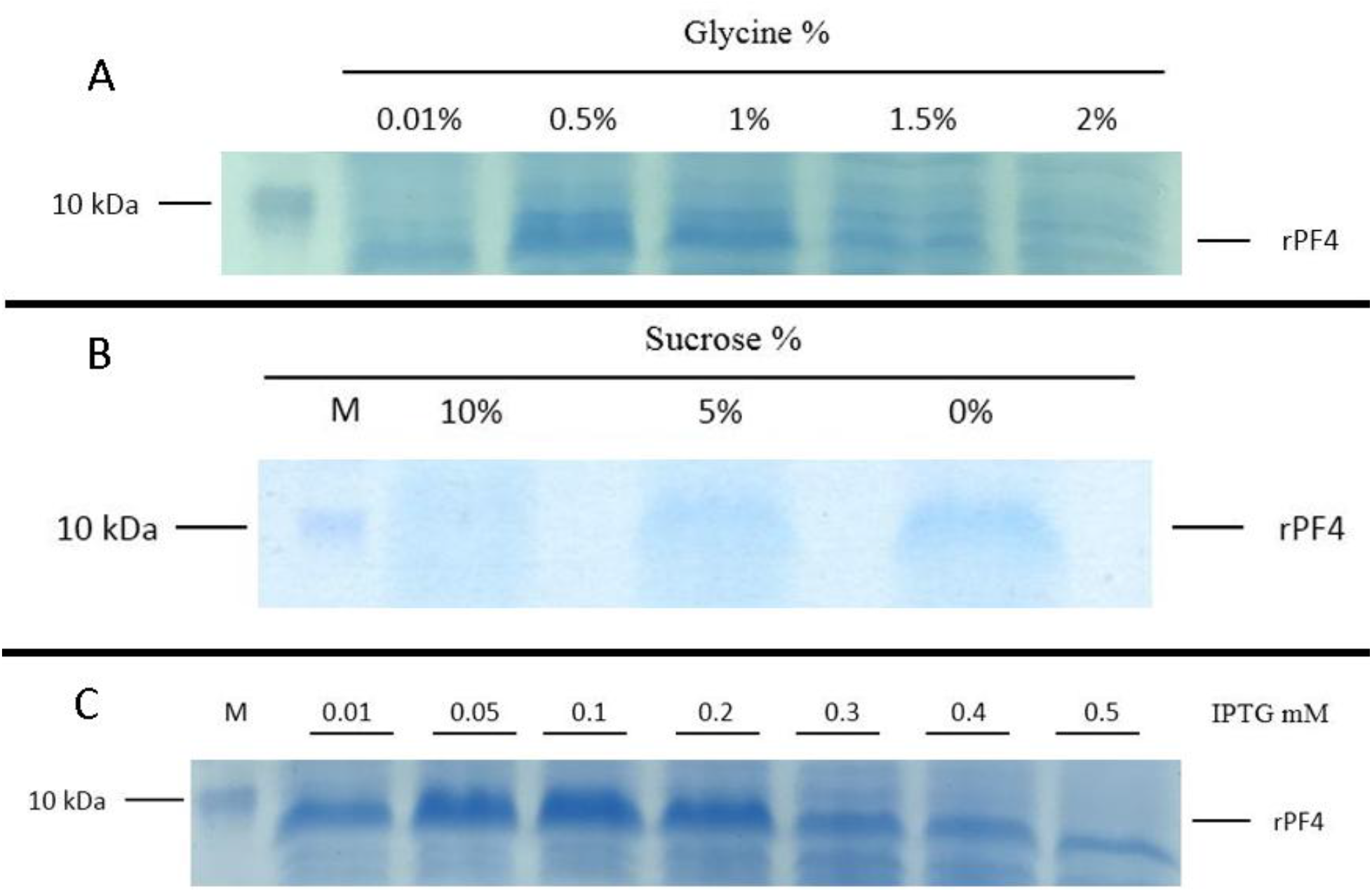
A) The effect of Glycine on protein secretion. Glycine supplementation facilitated the release of rPF4 into the extracellular space. Secretion at 0.5% glycine concentration demonstrated the highest efficiency. B) The effect of Sucrose on protein secretion. As opposed to Glycine, Sucrose supplementation negatively affected the secretion of rPF4. C) Fine tuning the IPTG concentration used to induce protein secretion. The induction stringency imposed on bacteria was changed from 0.01 through 0.5 mM concentration of IPTG, and 0.1 mM concentration of IPTG was found to be the optimum concentration to result in the best secretion level at 0.1 mM.

### 2.6 The trend of protein secretion over time

Total protein secretion per bacterial density was observed to increase over time during the first 22 hours but started to decline from then on (Sup fig 1).

### 2.7 a minor scale up in T2SS mediated protein secretion and purification of rPF4 in *E. coli*

Bearing in mind that cost-effectiveness along with achieving the highest yield is an important criterion to meet when dealing with recombinant protein production, a minor scale up to 40 ml devoid of any cultural supplementation was carried out. As mentioned previously induction under low IPTG concentration (0.1 mM) was performed. The secretion process continued for 48 hours and the aqueous media was subjected to protein purification. Proteins were eluted in 2ml fractions, and hereby, over 12 mg of rPF4 was purified from the media under native conditions (Fig. 10).

**Fig 10.**
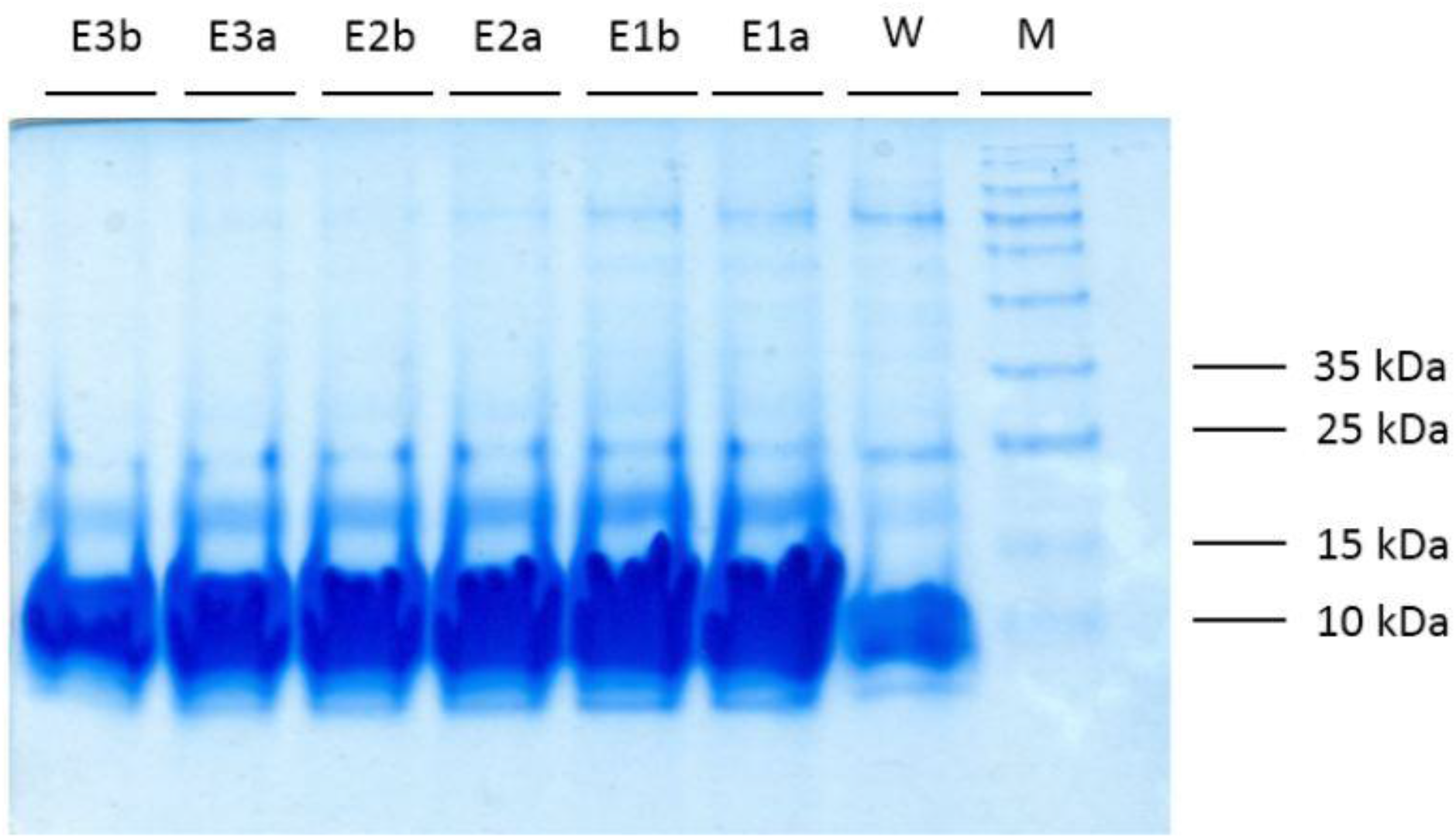
A minor scale up of T2SS-mediated protein secretion. The bacteria harboring the T2SS construct were grown in 40 ml of TB medium, induced and their supernatant was subjected to purification. Protein secretion was triggered at 0.1 mM IPTG and incubated for 48 hours. The column was washed with 10 ml of the washing solution containing 10 mM Imidazole. then, each time proteins were eluted with 2 ml elution buffer and ere collected in two 1 ml fractions, designated as “a” and “b”.

### 2.8 The synergistic effects of IPTG, Glycine and Triton X-100 on the improvement of protein secretion

A fractional factorial experiment was devised to examine the synergistic effect of various amount of IPTG, glycine, and Triton X-100 on the level of rPF4 secretion (Table 1). To facilitate execution of the experiment, sucrose was excluded from the analysis as the primary results conveyed negative effects. Since the secretion rate was negatively influenced by higher concentrations of IPTG, fractions of 0.3, 0.4 and 0.5 mM of IPTG were eliminated from the experiment, Furthermore, to reduce the detrimental effect of Triton X-100, the effect of lower concentrations were investigated. Hereby, protein secretion triggered at 0.05 mM IPTG with 1% glycine supplementation approximately yielded 286 μg/ml rPF4 secretion rate. Additionally, protein secretion triggered at 0.1 mM IPTG with 0.25% Triton X-100 supplementation approximately resulted in 700 μg/ml secretion rate. Nonetheless, co-supplementation of various amounts of glycine and Triton X-100 reflect an asymmetrical behavior. (Fig. 11, 12).

**Table 1.** Extracellular production of recombinant rPF4 (μg/ml) The bacterial cells were cultivated in TB medium harboring various concentrations of Glycine, Triton X-100 and IPTG. Protein secretion was continued for 48 hours.

| Triton X-100 | Glycine | IPTG mM |  |  |  |
| --- | --- | --- | --- | --- | --- |
|  |  | 0.01 | 0.05 | 0.1 | 0.2 |
| 0% | 0% | 83.2 | 149.57 | 122.51 | 61 |
|  | 0.5% | 99.17 | 102.75 | 97.61 | 137.5 |
|  | 1% | 224.7 | 286.14 | 262.21 | 97.14 |
| 0.25% | 0% | 356.15 | 614.97 | 701.75 | 412.25 |
|  | 0.5% | 320.37 | 425.95 | 286.78 | 238.92 |
|  | 1% | 305.38 | 311.64 | 245.28 | 122.11 |
| 0.5% | 0% | 78.24 | 93.92 | 3.34 | 108.53 |
|  | 0.5% | 49.42 | 57.52 | 233.39 | 100.12 |
|  | 1% | 139.42 | 86.18 | 70.18 | 4.40 |

**Fig 11.**
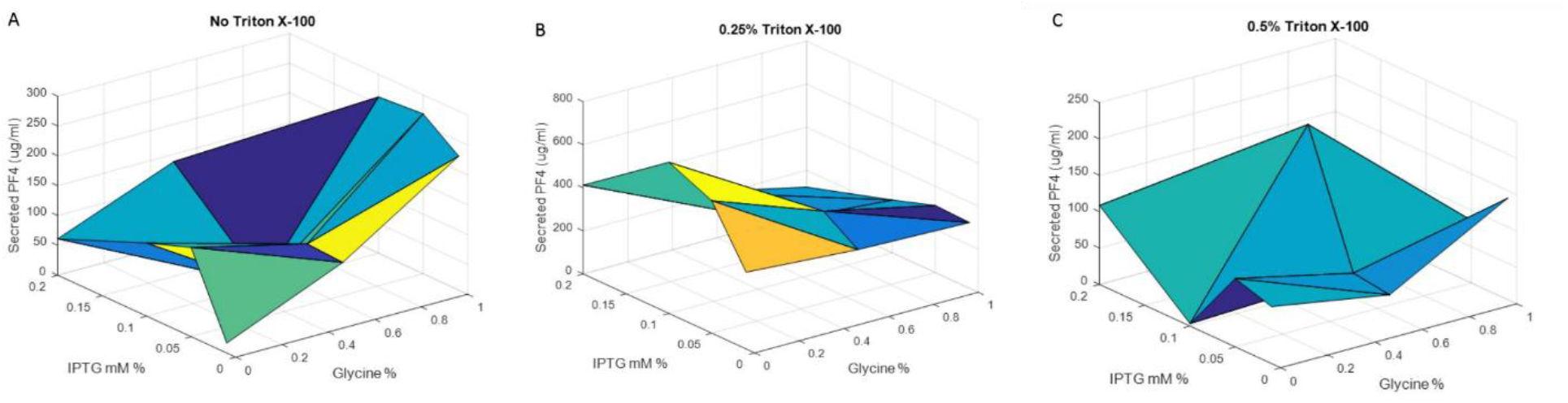
A demographic presentation of the synergistic effect on rPF4 secretion of various concentrations of Glycine, Triton X-100 and IPTG. A) In the absence of Triton X-100, the best secretion level is achieved as we move toward higher glycine concentrations and lower IPTG concentration between 0.1 and 0.15 mM. B) When the cocktail medium is supplemented with 0.25 % Triton X-100, the optimum secretion point is achieved by moving toward zero glycine concentration and 0.1mM IPTG. C) At 0.5 % Triton X-100 concentration, the optimum secretion level is brought about at midpoint glycine concentration and 0.1 mM IPTG.

**Fig 12.**
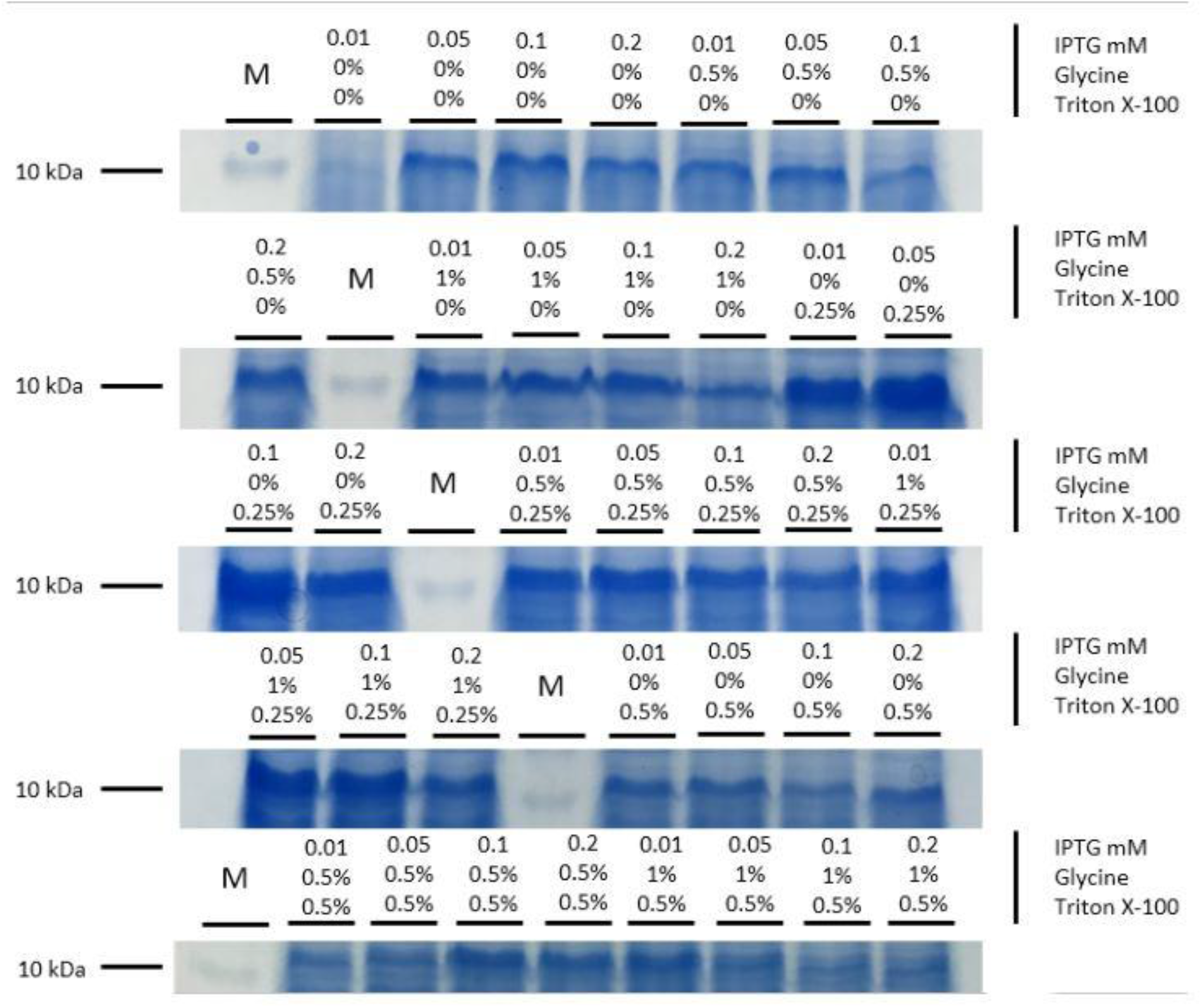
SDS-PAGE analysis of the synergistic effect of Glycine, Triton X-100, and IPTG. A combination of 36 different concentration cocktails of Glycine, Triton X-100, and IPTG revealed a synergistic effect between 0.5 % glycine, 0.5% Triton X-100, and 0.1 mM IPTG increases protein secretion approximately up to 233 μg/ml. however, a combination of 0.25 % Triton X-100 and 0.1 mM IPTG in the absence of glycine resulted in an approximately 700 μg/ml of secreted rPF4 as the optimum level.

### 2.9 Determination of protein concentrations using Image J software

As stated previously, of rising concentrations of BSA protein ranging from 100 to 1000 μg/ml were subjected to electrophoresis and the density corresponding to each band were determined accordingly the rPF4 protein concentrations were determined. To evaluate the relationship between the BSA concentrations and their corresponding density, the plots were analyzed by R squared value or the coefficient of determination which revealed a 97% correlation.

### 2.10 Recombinant rPF4 oligomerization, rPF4-Heparin ultra large complex formations, and zeta potential analysis

The Size distribution of complexes formed between rPF4 and heparin was investigated by Dynamic light scattering (DLS). DLS is governed by the Browning motion of particles in which the size of particles are inversely related. In the absence of heparin, the rPF4 at concentrations of 100, 200, and 400 μg/ml formed small particles of approximately 10 nm in size which is correlated to tetrameric oligomerization of the protein. Larger complexes of approximately 100 to 1200 nm in size were begin to form upon 5 to 20 unit commencement of unfractionated heparin to different concentrations of rPF4 which were further incubated for 15 to 120 minutes. Furthermore, Over-night incubation of rPF4 at 200 μg/ml concentration with 5 unit UFH produced larger complexes of roughly 600 and 1200 nm of size (Table 2) (Sup fig 2-11). Zeta analysis of 600 μg/ml of PF4 confirmed a positive charge of approximately 98 mV (Sup fig 12, 13).

**Table 2.** rPF4 tetramerization and heparin-rPF4 complex formation and oligomerization. When purified rPF4 was subjected to DLS, the size of particles detected span a range 8-14 nm in size therefore confirming tetramer formation by rPF4 secreted into the extracellular space. When rPF4 was mixed with heparin, oligomeric structures ranging in size from 100 to 1200 nm were detected indicating the tetrameric rPF4 proteins have acquired functional quaternary structures. The size of oligomeric structures increase in size in commensurate with an increase in incubation time and temperature up to 37 °C.

| **rPF4**<br>**($\mu\text{g/ml}$)** | **UFH**<br>**(unit)** | **Temperature** | **Incubation**<br>**Time (min)** | **Size**<br>**(nm)** | **SD** |
| --- | --- | --- | --- | --- | --- |
| 100 | - | 25 | - | 8.7 | $\pm 0.3$ |
| 100 | - | 25 | - | 14.5 | $\pm 0.8$ |
| 100 | 20 | 25 | 120 | 132 | $\pm 8$ |
| 100 | 20 | 25 | 120 | 86 | $\pm 3.4$ |
| 200 | - | 25 | - | 8.9 | $\pm 3.1$ |
| 200 | - | 25 | - | 12.8 | $\pm 5.2$ |
| 200 | 5 | 37 | Over night | 643 | $\pm 68.6$ |
| 200 | 5 | 37 | Over night | 1260 | $\pm 106.7$ |
| 400 | - | 25 | - | 7.8 | $\pm 3.4$ |
| 400 | 5 | 25 | 15 | 90 | $\pm 28.6$ |
| **400** | 10 | 25 | 15 | 126 | $\pm 47$ |

### 2.11 Raman spectroscopy analysis of rPF4

To ascertain that the recombinant rPF4 proteins are secreted in their native conformation, 2 mg of rPF4 was freeze dried and subjected to Raman spectroscopy. The spectrum was analyzed in different Raman shifts to identify formations of disulfide bonds and secondary structure of rPF4. Ensuing decomposition and a linear base line subtraction in amid I region, rPF4 was unveiled to be approximately comprised of 43.5% Random coil, 32.5% β-sheet, 18.6% α-helix, and 4.9% turn, which is in concordance with the crystal structure PF4 reported by the protein data bank. The native PF4 structure comprised of two disulfide bonds with pivotal roles in native functional conformation of the protein and by careful analysis of the Raman spectrum the formation these two critical disulfide bonds was revealed (Fig. 13).

**Fig 13.**
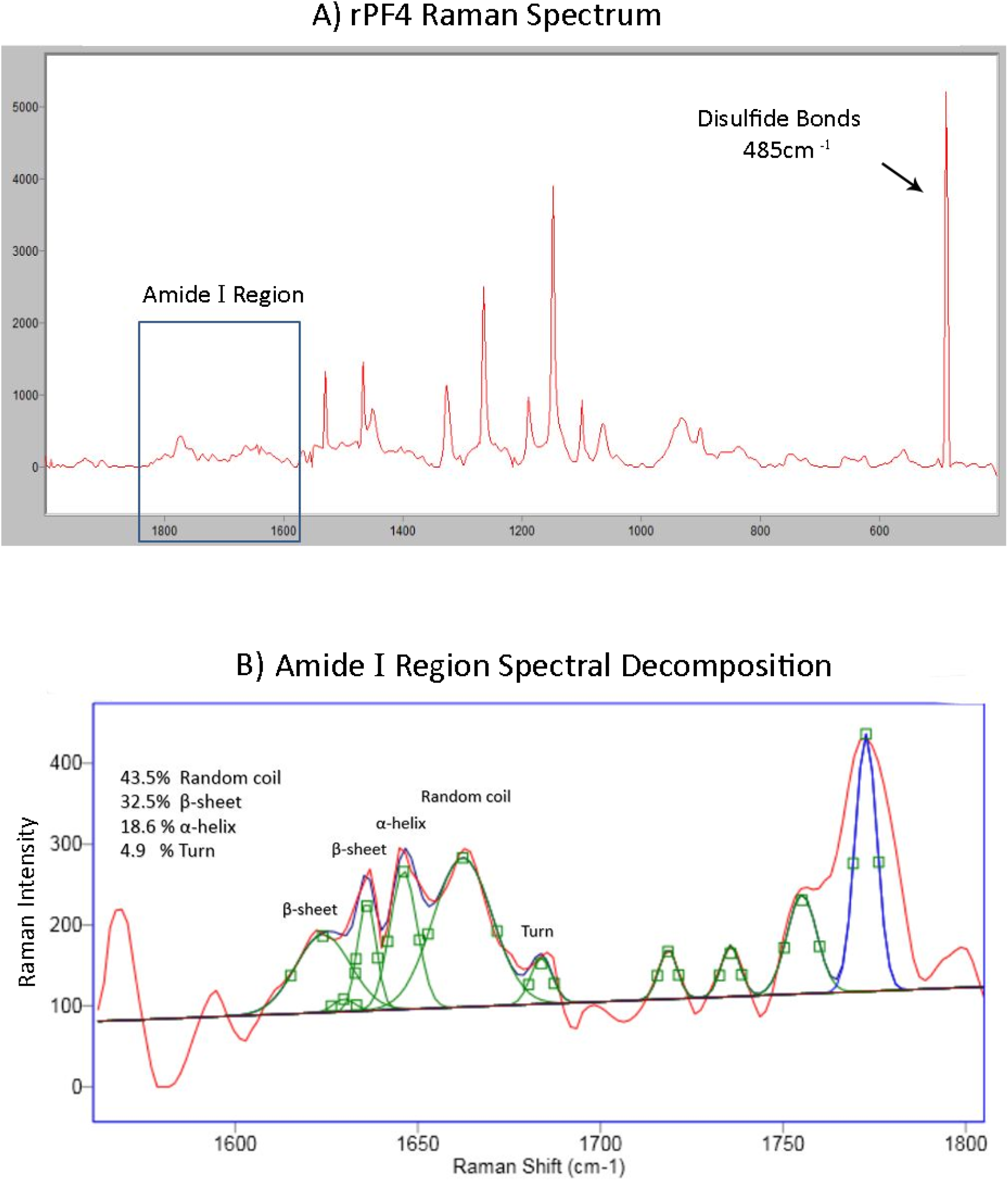
Recombinant rPF4 secondary structure and disulfide bond formations. By exploiting Raman spectroscopy, the conformation of the secreted recombinant rPF4 was analyzed. (A) Represents disulfide bond formation at the 485 cm-1 of Raman shift. (B) Following decomposition of the amide I region, the secondary structure of the rPF4 was determined and revealed to be in perfect agreement with predictions based on x-ray crystallography data.

## 3. Discussion

*E. coli* has been the chief bacterial host for the production of recombinant proteins with commercial and medical significance. A kaleidoscopic variety of genetically engineered hosts, expression vectors, and expression strategies have made this process significantly convenient(22, 23). Intracellular protein expression has been in widespread use, albeit complicated by hurdles including inclusion bodies(24), protein degradation due to the presence of unleashed proteases, and liberation of lipopolysaccharide (LPS), demanding cumbersome downstream procedures such as denaturation and refolding(25), endotoxin removal(26, 27) and successive steps of protein purification.(28).

Extracellular protein production seems to be promising as it has the potential to circumvent the need for breaking the cells open, enhances protein purity and paves the way for the large-scale industrial protein production(13). We envisioned that the T1SS and the T2SS secretory pathways of *E. coli* would be suitable for rPF4 production. Several factors contribute to the superiority of recombinant protein secretion in *E.coli* over the conventional cytoplasmic expression. Target proteins are sequestered away from a variety of destructive proteases as they are protected by the secretory machinery. SecB, as a pivotal component of the type II secretory pathway, grabs onto and holds the target protein in a semi-folded conformation thus retaining their competency for the post-translational translocation through the inner membrane into the periplasmic space by preventing protein aggregation(29). Based on previous studies indicating the dependency of pelB signal sequence on secB to direct protein secretion(18), we speculate that coexpression of secB may also improve the secretion rate of rPF4 via type II secretory pathway by augmenting T2SS machinery through providing an excessive number of secB proteins, compensating for the rate-limiting amount of secB at disposal. Furthermore, based on our observations, in the case of toxic recombinant proteins, secB sequesters the target proteins and diminishes their toxicity to a large extent (data not published). As opposed to the highly crowded intracellular area where the mean distance between the proteins is less than the size of an average protein which results in a huge thermodynamic driving force for protein aggregation (30), the ample extracellular space provides the exported recombinant proteins with enough room to accumulate and remain in their native state.

However, although a considerable amount of rPF4 was observed to be secreted into the extracellular space, we envisioned that the periplasmic environment is still teeming with large quantities of rPF4, and to release the proteins trapped there while continuing to preserve the integrity of the outer membrane and minimizing the risk of LPS contamination to the minimum, small amounts of Glycine and Triton X-100 supplementation were included, which revealed to be highly effective perhaps by slightly increasing the outer membrane permeability. Furthermore, as IPTG triggers the protein expression through de-repressing the T7 promoter, we sought to evaluate the optimum depression allowing the components of the secretory apparatus to keep up with the pace of protein production so as to achieve maximum protein secretion. At higher concentrations of IPTG the secretary system gets exhausted and fails to assist in efficient protein secretion, and as consequence lower secretion yield is observed. On the contrary, a very mild induction of protein expression highly favors protein folding process, solubility, and an efficient transportation of proteins into the extracellular milieu.

A higher proportion of chaperones residing in the periplasmic environment(29) creates a more favorable compartment for protein folding and correct disulfide bond formation, particularly when dealing with sophisticated eukaryotic proteins with more complicated folding kinetics. Therefore, by exporting the recombinant proteins into the periplasm there is a higher likelihood of assuming the native functional state by the recombinant proteins. From a different perspective, it is notable that the lower levels of proteases in the periplasm boosts protein stability by avoiding unwanted degradation(13). Furthermore, Cleavage by the signal peptidase (for instance the Lep protease) of the N-terminal signal sequence leads to an authentic N-terminus by rendering the target protein devoid of the initial Methionine, restriction enzyme scar sites and other unwanted junk sequences that may adversely affect the native protein structure and function(31).

A variety of measures were taken to mitigate the shortcomings of cytoplasmic recombinant PF4 production to achieve enhanced solubility, higher yields, robust folding, efficient oligomerization, and increased purity through cumbersome manipulation of intracellular expression systems and successive steps of technically demanding protein purification strategies(32, 33). However, extracellular recombinant PF4 production was found to alleviate many of the aforementioned downsides compared to intracellular protein expression. Ultra-large complex formation between unfractionated heparin and secreted rPF4 revealed a high protein solubility, proper folding, and efficient tetrameric oligomerization, as confirmed by Dynamic light scattering and Raman spectroscopy. The secretion of recombinant rPF4 to the extracellular medium also circumvents the arduous task of removing LPS as it preserves the bacterial cell integrity, since it avoids the need to rupture bacterial cell wall. Proper disulfide bond formation has always been a major concern in cytoplasmic protein expression methods, which commonly demands difficult downstream procedures, however, by harnessing the secretory apparatus of the *E. coli* as the workhorse in molecular biology and biotechnology, these conundrums are eliminated.

Based on the quantity of protein mass accumulated in the bacterial culture, the classical HylA T1SS of *E. coli* directly ferrying recombinant rPF4 proteins to the extracellular milieu was found to be inferior to the SecB-dependent T2SS. The low expression levels and early exhaustion of molecular constituents involved in classical HylA T1SS might be the rate-limiting agents and thus accounted for the inefficiency of rPF4 secretion using this system. It is envisaged that an all-in-one vector furnishing a surplus of these restrictive components might boost the system’s capacity for a more efficient T1SS-based export of recombinant PF4 outside the cell.

Here Ni-NTA purification system was employed but shifting to an alternative approach including heparin affinity chromatography circumvents the need to add a patch of extra Histidine amino acids leading to exact structural similarity with the native platelet-derived factor.

## 4. Conclusion

In conclusion, employing an extracellular protein production approach facilitates PF4 production, purification, and biochemical analysis to a large extent which remedies the many shortcomings associated with conventional cytoplasmic expression of recombinant proteins and promises a more efficient and cost-effective approach for future industrial applications.

## 5 Materials and methods

### 5.1 Strains and reagents

The restriction enzymes (SlaI, SalI, and NdeI) were obtained from Jena Bioscience (Jena, Germany), T4 DNA ligase was purchased from Thermo Fisher Scientific (Grand Island, NY). Anti-His Tag antibody (Abcam Company, Cambridge, UK) was used to perform western blotting method. Immobilized metal affinity chromatography (IMAC) using Ni-NTA matrix (Qiagen) was employed for purification purposes. The synthetic construct was ordered from Biomatik Company (Biomatik, Ontario, Canada). To alleviate problems regarding *E. coli* codon bias Biomatik’s proprietary codon optimization service was exploited. The agarose gel and plasmid extraction kits were purchased from Bioneer (South Korea). Isopropyl beta-D-thiogalactopyranoside (IPTG) and Kanamycin sulfate were acquired from Bio-Basic (Markham, Canada) and Merk (Germany) respectively. pET26b was utilized as the expression vector. BL21 (DE3) and DH5a strains of *E. coli* were acquired from Diagnostic Laboratory Sciences and Technology Research Center, Shiraz, Iran. The *E. coli* DH5a was employed for restriction digestions and plasmid extraction purposes. The *E. coli* BL21 (DE3) was exploited as an expression host.

### 5.2 DNA constructs

The human PF4 sequence (**accession Number: P02776**) and the *E. coli* hemolysin alpha (**accession Number: P08715**) signal sequence (HlyAs) which corresponds to residues 965-1024 of α-hemolysin with the NdeI and SlaI restriction sites at the 5’ and 3’ termini and a SalI restriction site prior to HlyAs, was commercially custom-synthesized and cloned into pET26b at the NcoI and XhoI sites in the multiple cloning site of the vector. As part of the pET26b vector multiple cloning sites, a pelB signal sequence is placed upstream of the PF4 coding sequence. To obtain the constructs to be exported using type I and II secretion systems, the synthetic 3 in one construct harboring the desired signal sequences was subjected to a series of digestions and ligations. the pET26b-pelB-rPF4 construct was made through sequential digestions using SalI and SlaI enzymes whereas pET26b-rPF4-HlyAs construct was obtained through single NdeI digestion, Moreover, the pET26b-rPF4 construct was generated through a single digestion of pET26b-pelB-rPF4 construct by the NdeI enzyme. As the HlyAs signal sequence needs to be free at the c-terminal end of the protein for its proper functioning, a 6-His patch was included between the PF4 gene and HlyAs through a linker sequence harboring a Tev recognition site. Located downstream of the HlyAs is a stop codon followed by another 6-His patch and a second stop codon. After removal of the HlyAs signal, the first stop codon is excised out allowing the second His patch to be placed downstream of the PF4 gene, ensued by the second stop codon enabling the expression of pelB-rPF4-His to halt (Fig. 1). The ligation reactions were performed by T4 DNA ligase. All the reactions were carried out according to the manufacturers’ protocol. Colony PCR assay, and nucleotide sequencing methods were employed to confirm the correct plasmid constructions.

### 5.3 Expression of the pelB-rPF4, rPF4-HLA and rPF4 proteins

A single colony from the bacteria transformed with the previously prepared constructs was grown into 5 ml of Luria-Bertani broth (LB) separately, each containing 70 μg/ml Kanamycin and incubated overnight at 37 °C, 130 rpm. One hundred microliters of the starter cultures was transferred into 7 mL of fresh LB or TB (Terrific Broth) media containing 70 μg/mL Kanamycin. Based on our previous experience with low molecular weight (MW) protein expression, expression was triggered at higher IPTG concentrations and accompanied by a short induction duration. The cells were grown at 37 °C, 150 rpm until reaching 0.6-1.0 (OD 600) concentration. Then, protein expression was induced using 2 mM IPTG at 37 °C, 200 rpm for 2 h. The cells were harvested by centrifugation at 5000g for 15 min at room temperature.

### 5.4 Verification of protein expression using SDS-PAGE

The expression of recombinant proteins in control and induced samples were analyzed employing SDS-PAGE. The harvested The bacterial cells, were dissolved in denaturing lysis buffer (NaH_2_PO_4_ 100 mM, Tris-HCl 10 mM, urea 8 M, and pH 8.0) and broken open using sonication. The resulting crude bacterial extract was centrifuged at 10000 g for 30 min at 4 C. Twenty microliters of the cleared supernatants corresponding to rPF4, pelB-rPF4 or rPF4-HLAs was mixed with 20 μl of Laemmli buffer and boiled at 100 C for 5 min. The samples were loaded onto 13.5% (v/v) resolving, 5% (v/v) stacking gels. A run at 80 V for 35 min stacked the proteins and further ensued by 150 V for 70 min to separate the proteins. The gel was stained using Coomassie Brilliant Blue and the rPF4-HlAs (16 kDa), pelB-rPF4 (11.4 kDa), and rPF4 (8.8 kDa) proteins were detected.

### 5.5 Purification of the recombinant rPF4 protein

To further validate the rPF4 production, the proteins were purified employing immobilized metal affinity chromatography (IMAC) method using Ni-NTA matrix. Cleared lysate was prepared as described in section 5.4. Lysis buffer contained 10 mM Imidazole and 0.5 M NaCl to decrease the chance of unspecific protein bindings to the matrix. Binding step lasted for 2 hours with 200 rpm shaking at the ambient temperature. The column was washed 2 times with the wash buffer (NaH_2_PO_4_ 100 mM, Tris-HCl 10 mM, imidazole 10 mM, NaCl 0.5 M, Urea 8 M, pH 8.0) and four 0.5 ml eluates were collected using elution buffer (NaH_2_PO_4_ 100 mM, Tris-HCl 10 mM, imidazole 250 mM, NaCl 0.5 M, Urea 8 M, pH 8.0).

### 5.6 rPF4 protein secretion mediated by type I and II of gram negative bacterial secretory pathways

rPF4 secretion mediated by type I and type II gram-negative bacterial secretory pathway was analyzed. A two set of single colony *E. coli* BL21 harboring either pET26b-pelB-rPF4 or pET26b-rPF4-HLAs plasmids were grown into 5 ml Luria Bertani broth (LB) containing 70 mg/ml Kanamycin and incubated overnight at 37 °C, 130 rpm. One hundred microliters of the starter culture was transferred into 7 mL of fresh LB and TB media containing 70 mg/mL Kanamycin. The cells were grown at 37 °C, 150 rpm until reaching 0.6-1.0 (OD 600) concentration, subsequently protein expression was triggered by 0.5mM IPTG at 29 °C.

### 5.7 SDS-PAGE analysis of purified extracellular milieu from *E. coli* BL21 (DE3) cells producing recombinant pelB-rPF4 and rPF4-HLAs and rPF4 proteins

To analyze type I- and II-mediated secretion, a single *E. coli* BL21 colony containing either pET26b-pelB-rPF4, pET26b-rPF4-HLAs or pET26b-rPF4 was grown into 5 ml LB containing 70 μg/ml Kanamycin and incubated overnight at 37 °C, 130 rpm. One hundred microliters of either starter cultures were transferred into 10 ml of fresh LB or TB media containing 70 μg/ml Kanamycin. The cells were grown at 37 °C, 150 rpm until reaching 0.6-1.0 (OD 600) concentration and protein expression was induced using 0.5 mM IPTG at 29 °C, 200 rpm for up to 24 h. Subsequently, the TB and LB media expected to contain the secreted recombinant rPF4 protein was subjected to purification using immobilized metal affinity chromatography (IMAC) under native conditions. SDS-PAGE was employed to analyze the secreted rPF4 protein purified from the extracellular milieu.

### 5.8 Verification of protein secretion using Western blotting

To further validate rPF4 protein expression and secretion we inspected the aqueous media for the presence of recombinant proteins. Proteins were verified by SDS-PAGE and western blotting methods. Electrophoresis was performed on 13.5% gel according to section 5.4. Western blotting was performed using specific antibody. The protein bands were effectively blotted to PVDF membrane using the semidry protocol mediated by Towbin transfer buffer at 20 V for 70 min. The membrane was blocked in a 20ml blocking buffer containing 1% (w/v) skim milk for 1 h on a rocking platform and rinsed in 30 ml of TBST wash buffer containing 0.02% (v /v) tween. The membrane was immersed in horseradish peroxidase-conjugated antibody diluted in blocking buffer (1:1000 dilution) for 60 min on a rocking platform at room temperature. The membrane was washed 6 times in TBST (containing 0.02% (v/v) Tween-20). Finally, the protein bands of interest were visualized using diaminobenzidine (DAB) solution as the enzymes substrate.

### 5.9 Investigating protein secretion level utilizing the effect of IPTG, Triton X-100, Glycine and sucrose individually

A single colony of *E. coli* BL21 (DE3) containing pET26b-pelB-rPF4 was inoculated into 5 ml LB containing 70 μg/ml Kanamycin and grown overnight at 37 °C, 130 rpm. One hundred microliters of the starter cultures was transferred into four groups of 100 ml of fresh TB media containing 70 μg/ml Kanamycin. The cells were grown at 37 °C, 150 rpm until reaching 0.6-1.0 (OD 600) concentration. The culture was divided into 10 ml flasks to individually analyze the effect of IPTG (0.01, 0.05, 0.1, 0.2, 0.3, 0.4 0.5 mM), glycine (0%, 0.5%, 1%, 1.5%, 2%, 3%, 4%, 5% w/v), Triton X-100 (0%, 1%, 2%, 3% w/v) and sucrose (0%, 5%, 10% w/v). The cocktail media containing glycine, sucrose and Triton X-100 supplementations, cultures were supplied with IPTG to the final concentration of 0.5 mM.

### 5.10 Investigating the trend of proteins secretion over time

To investigate the optimum secretion duration which results in the maximum level of protein secretion and to observe the trend of protein secretion over time, the protein content of the extracellular milieu was analyzed at several time points. To this end, 10 ml of induced culture harboring pET26b-pelB-rPF4 construct, was subjected to analysis 5.5, 14, 20, 26, 39 and 48 hours post-induction.

### 5.11 Investigating synergistic effect of IPTG, Triton X-100, and Glycine supplementations on protein secretion level

To further improve the secretion process, a fractional factorial experiment was employed to evaluate the effects of Triton X-100, Glycine and optimized concentration of IPTG. To this end, 400 mL of TB cultures of pelB-rPF4 containing 70 μg/ml kanamycin that reached the 0.6-1.0 (OD 600) of concentration, was divided into 36 flasks each containing 5 mL of culture. Next a combination of varied volumes of 100mM IPTG, 10% glycine, and 10% Triton X-100 were added to the shaking flasks until reaching the final concentrations of IPTG (0.01, 0.05, 0.1 and 0.2 mM), glycine (0%, 0.5%, 1% w/v), and Triton X-100 (0%, 025%, 0.5% w/v), respectively. The flasks were supplied with an appropriate volume of TB medium to adjust to the final volume of 10 ml per shaking flask, and the flasks were further cultivated at 29 °C with shaking (150 rpm) for 48 hours.

### 5.12 Protein secretion analysis method exploiting Image J software

SDS-PAGE was employed to analyze extracellular protein secretion. The density of protein bands was determined by employing Image J software. in order to equate The density of protein bands with their respective concentration a set of rising amount of Bovine serum albumin (BSA) protein ranging from 100 to 1000 μg/ml were subjected to electrophoresis and the density corresponding to each band were determined. Next, the density was plotted against the concentration of proteins which allows to approximately determine rPF4 secretion amount.

### 5.13 Analyzing rPF4 oligomerization, rPF4-heparin ultra-large complex formations and zeta potential measurements

rPF4 oligomerization and rPF4-heparin complex formations were studied using a dynamic light scattering (DLS) technique (Nanopartica SZ 100; HORIBA Ltd, Kyoto, Japan) with a fixed 173 scattering angle and a 633-nm helium-neon laser. Data were analyzed using Horiba SZ 100 for Windows [Z Type] software version 2.20 (Nanopartica SZ 100; HORIBA Ltd, Kyoto, Japan). Furthermore, Zeta potential experiments were performed, employing Horiba SZ 100.

### 5.14 Determining secreted rPF4 secondary structures by Raman spectroscopy

rPF4 protein was subjected to lyophilization and the resulting powder was utilized for structural investigation. The experiment was performed at room temperature having a relative humidity of 20% using a LABRAM HR Raman spectrometer (Horiba Jobin Yvon, Villeneuve d’Ascq, France) coupled to a confocal microscope. The laser light at 532 nm (INNOVA 70C Series Ion Laser, Coherent, Santa Clara, CA) was used to illuminate the samples, and recording was carried out in 20 second intervals. 100X objective lens was used to focus the laser beam on lyophilized protein samples (0.9 NA, Olympus, Melville, NY). Spectral analysis was implemented using GRAMS/AI™ software (GRAMS SUITE 9.2, Thermo Fisher Scientific Inc.) and a linear baseline subtraction was performed in the amide I region spanning the 1600–1700 cm-1 range.

## Declarations

### Abbreviations

PF4: Platelet Factor 4
T1SS: Type I secretion system
T2SS: Type II secretion system
LB: Luria-Bertani
TB: Terrific Broth
DLS: Dynamic Light Scattering

## Acknowledgments

Authors would like to thank the staff at the Diagnostic Laboratory Sciences and Technology Research Center (DLSTRC) for their support.

## Authors’ contributions

GT and MNT supervised the project, MNT and SA conceived, designed and carried out the experiments. MNT and SA analyzed the data, FT and FZ participated in analyzing the data and performing the experiments respectively, NA created the schematic figures, AB and AR provided useful discussions. GT, MNT and SA wrote the manuscript. MNT and SA critically revised the manuscript. Authors studied and approved the manuscript.

## Availability of data and materials

The data supporting the findings of this study are available within the paper and its additional file. Further informations are available from the corresponding authors on request

## Ethics approval and consent to participate

Not applicable

## Consent for publication

All authors read and approved the final manuscript.

