## Supplementary material for "High-yield Production of Recombinant Platelet Factor 4 Protein by Harnessing and Honing the Gram-negative Bacterial Secretory Apparatus": Sup

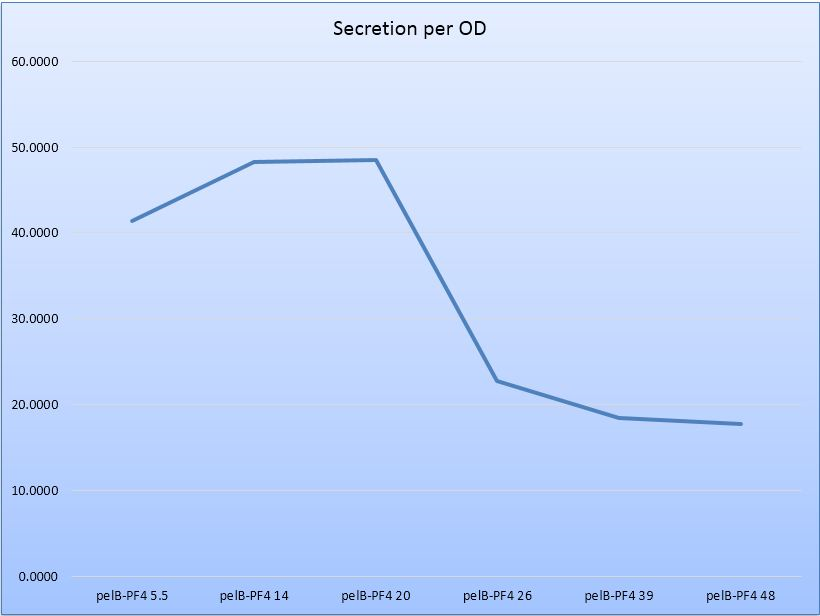

Supplementary Fig 1) the trend of protein secretion over time

Protein secretion per bacterial density follows an increasing trend for the first 22 hours, and begins to decline then after. The vertical axes, indicates secretion per bacterial density, in terms of micrograms per milliliter per OD.

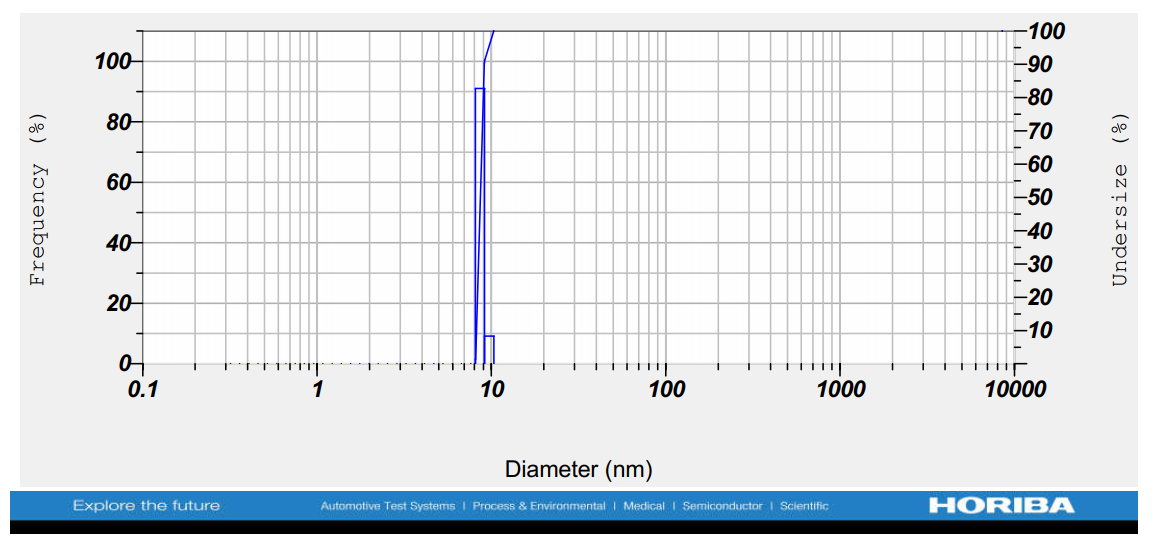

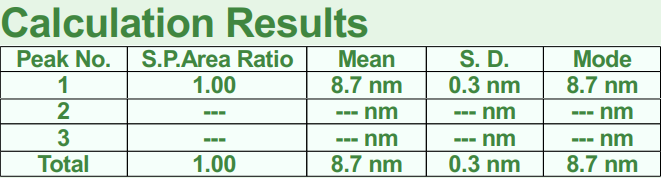

Supplementary Fig 2) 100 µg/ml rPF4 oligomerization analysis.

100 µg/ml of rPF4 was subjected to DLS measurements.

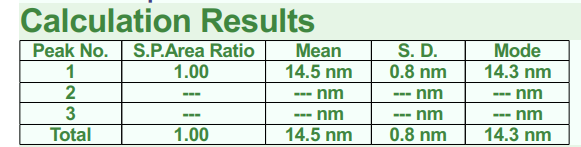

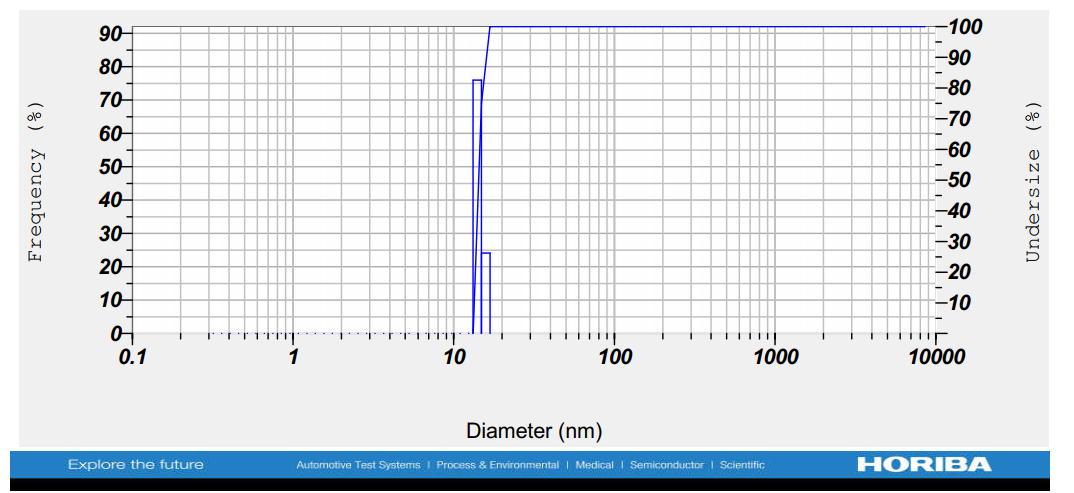

Supplementary Fig 3) 100 µg/ml rPF4 oligomerization analysis.

100 µg/ml of rPF4 was subjected to DLS measurements.

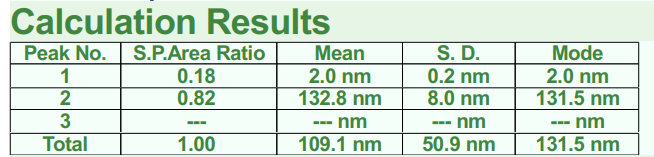

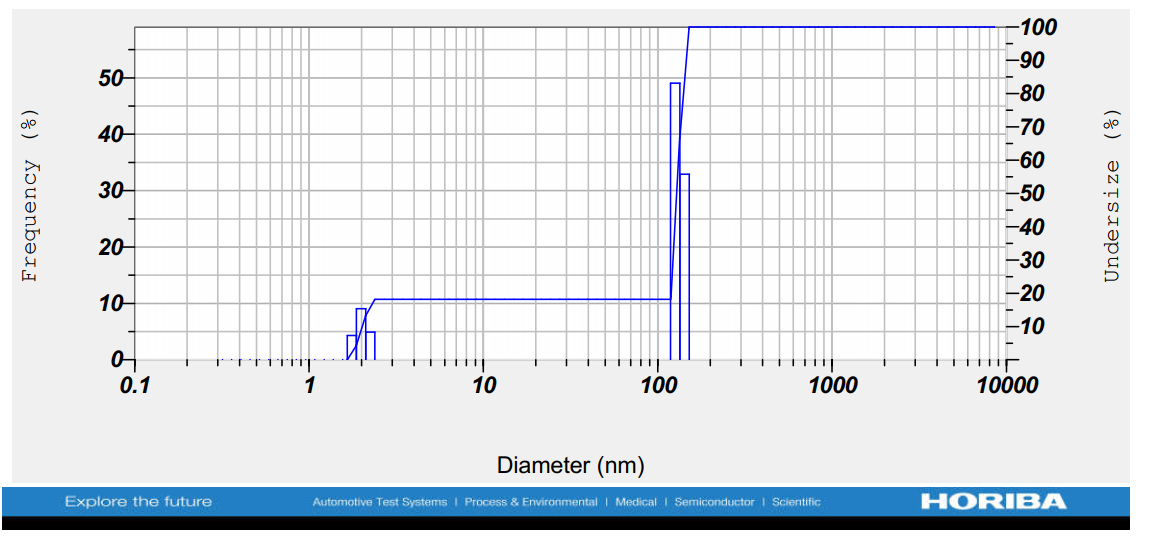

Supplementary Fig 4) rPF4-heparin complex formation analysis.

100 µg/ml of rPF4 was supplemented with 20 units/ml of UFH and was further incubated for 2 hours.

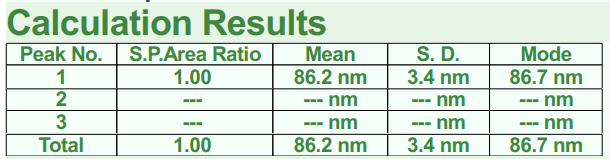

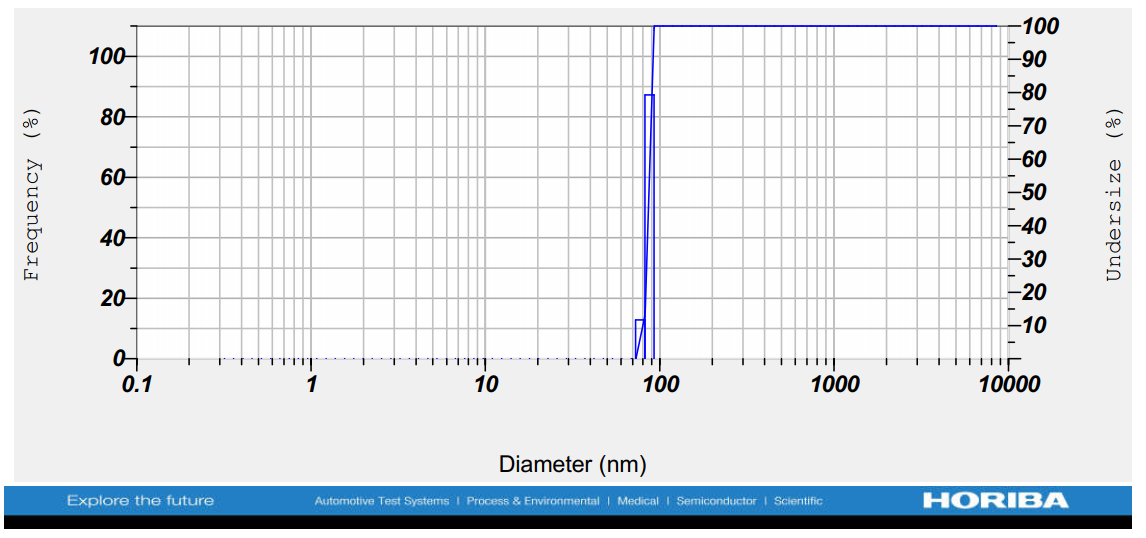

Supplementary Fig 5) rPF4-heparin complex formation analysis.

100 µg/ml of rPF4 was supplemented with 20 units/ml of UFH and was further incubated for 2 hours.

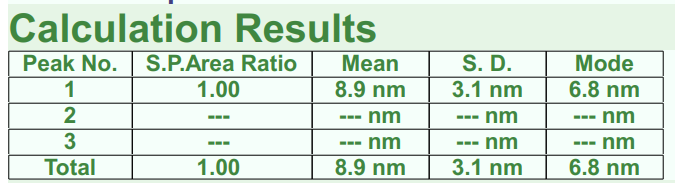

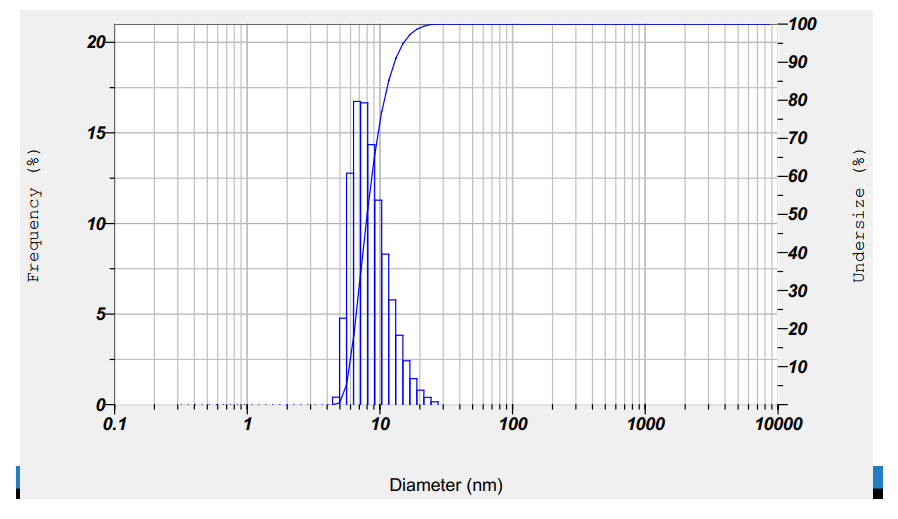

Supplementary Fig 6) 200 µg/ml rPF4 oligomerization analysis.

200 µg/ml of rPF4 was subjected to DLS measurements.

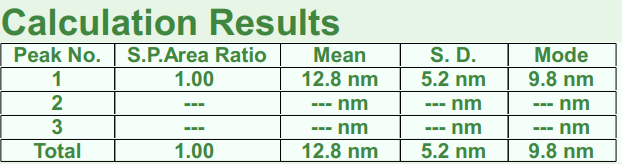

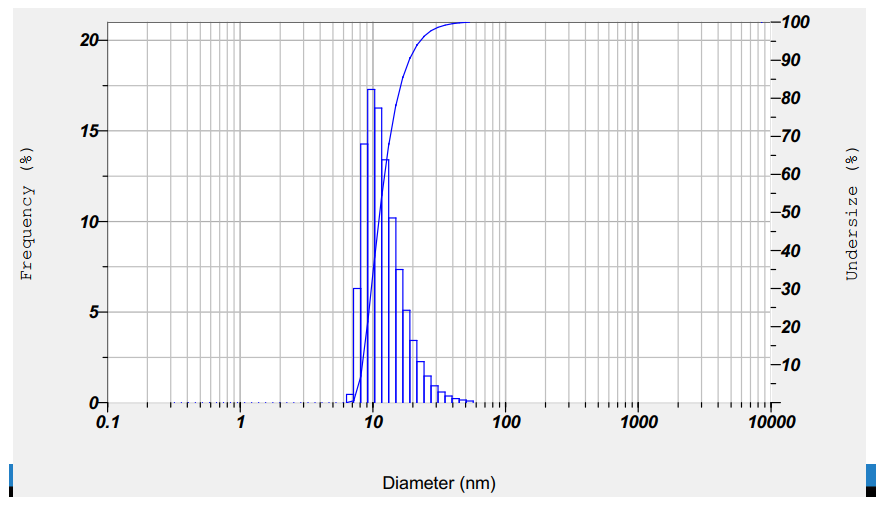

Supplementary Fig 7) 200 µg/ml rPF4 oligomerization analysis.

200 µg/ml of rPF4 was subjected to DLS measurements.

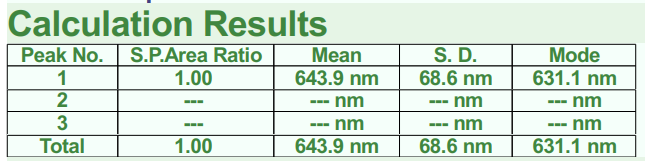

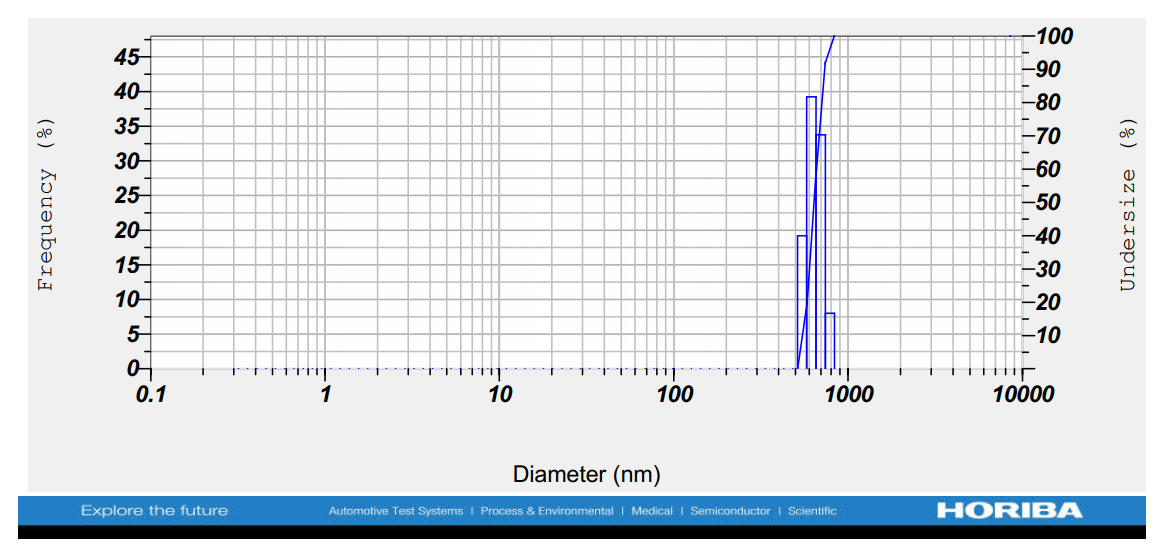

Supplementary Figure 8) rPF4-heparin complex formation analysis.

200 µg/ml of rPF4 was subjected to DLS measurements. 200 µg/ml of rPF4 was supplemented with 5 units/ml of UFH, and overnight incubation was further carried out.

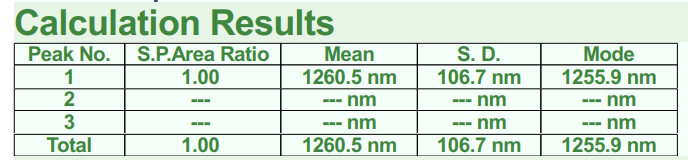

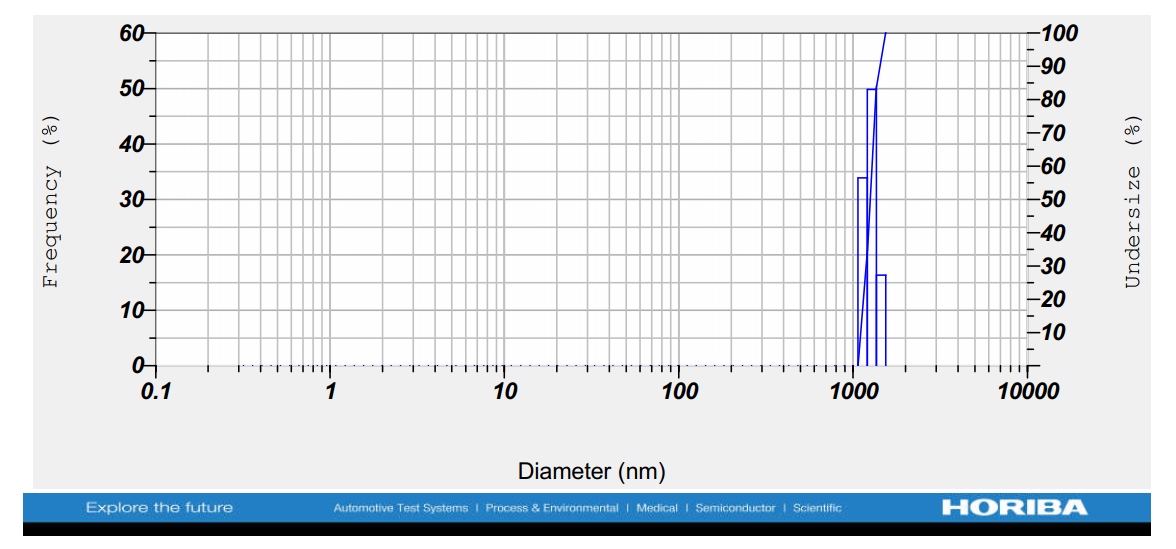

Supplementary Fig 9) rPF4-heparin complex formation analysis.

200 µg/ml of rPF4 was subjected to DLS measurements. 200 µg/ml of rPF4 was supplemented with 5 units/ml of UFH, and overnight incubation was further carried out.

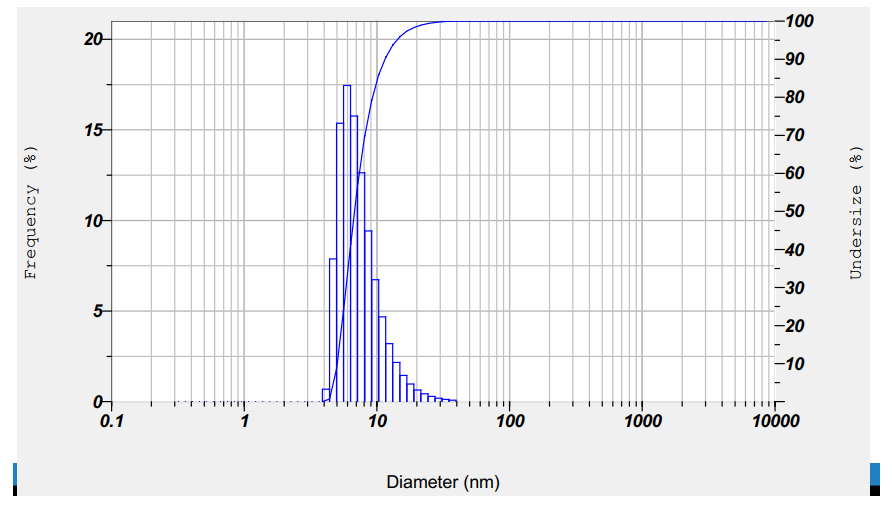

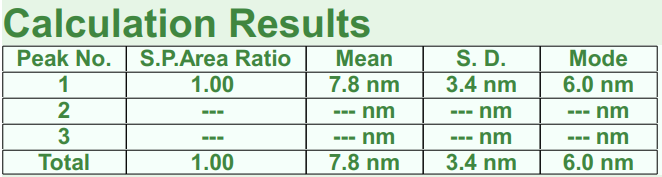

Supplementary Figure 10) 400µg/ml rPF4 oligomerization analysis.

400µg/ml of rPF4 was subjected to DLS measurements.

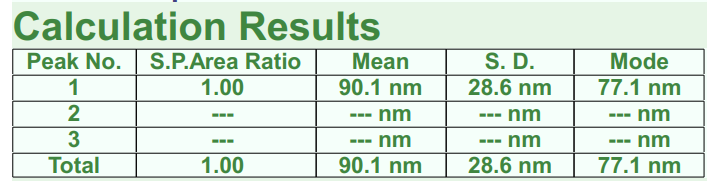

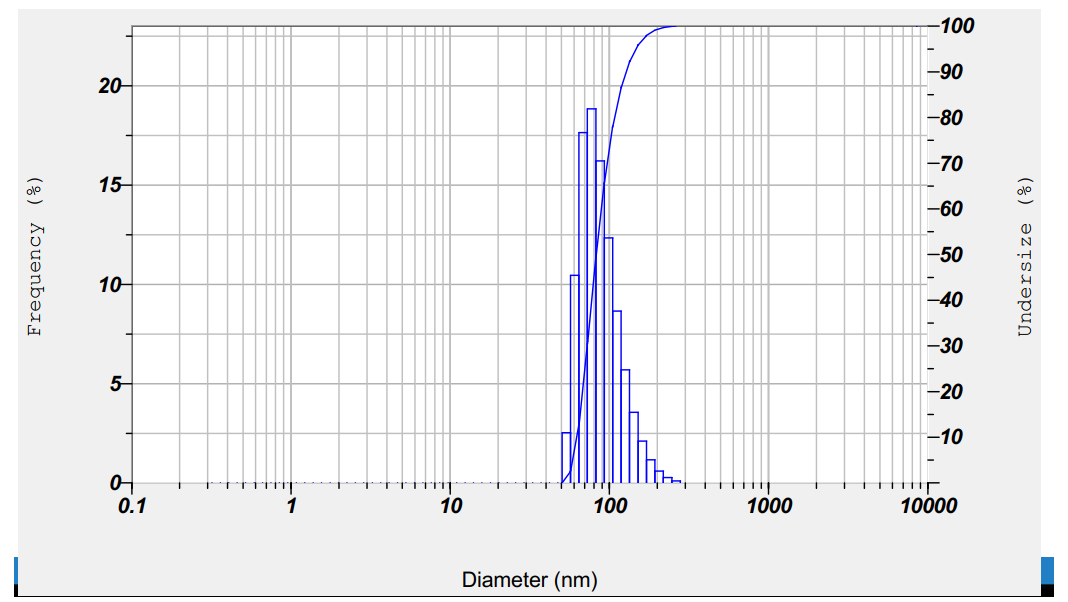

Supplementary Figure 11) rPF4-heparin formation analysis.

400 µg/ml of rPF4 was supplemented with 5 unit/ml of UFH, and was further incubated for 15 minutes.

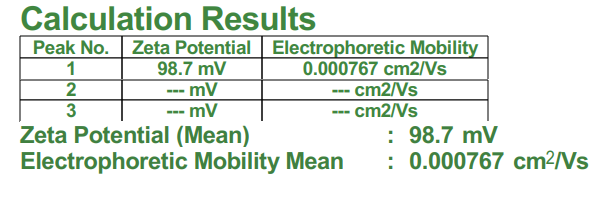

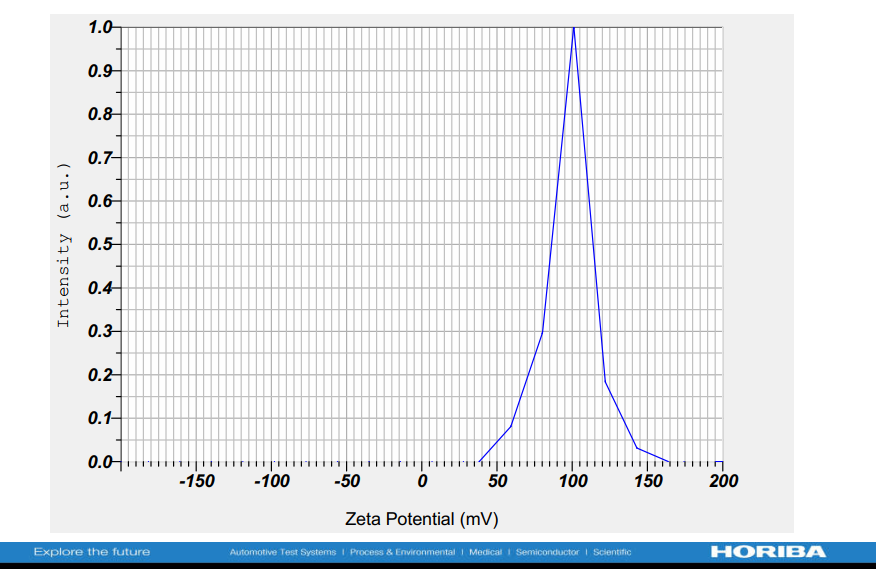

Supplementary Figure 12) rPF4 zeta potential analysis.

600 µg/ml of rPF4 was subjected to analysis

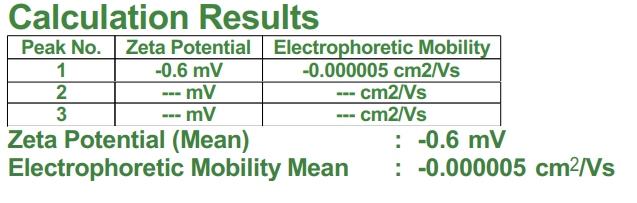

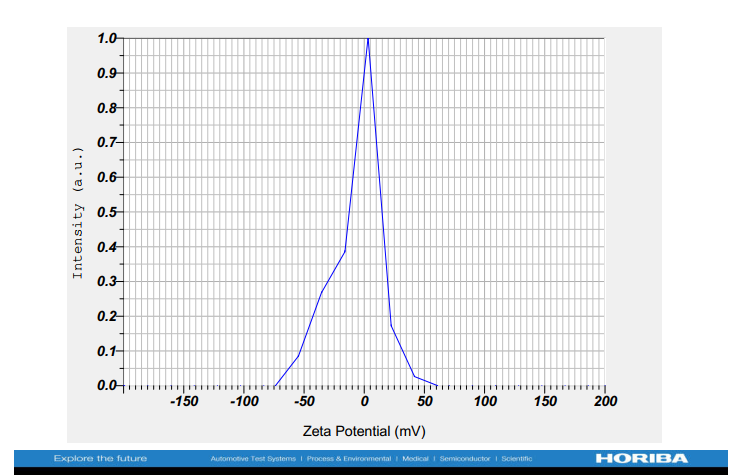

Supplementary Figure 13) Zeta potential analysis the elution buffer devoid of rPF4. 600 µg/ml of rPF4 was subjected to analysis
